# Loss of Cdk5rap2 triggers cellular senescence via β-catenin-mediated downregulation of WIP1

**DOI:** 10.1101/2020.07.09.194761

**Authors:** Xidi Wang, Patrick Sipila, Zizhen Si, Jesusa L. Rosales, Xu Gao, Ki-Young Lee

## Abstract

Loss-of-function mutations in Cdk5rap2 is associated with the developmental disorders, primary microcephaly and primordial dwarfism, but the underlying molecular link remains obscure. Here, we show that Cdk5rap2 loss in BJ-5ta human fibroblasts triggers senescence that is associated with proliferation defect, which is manifested as small body size in Cdk5rap2^*an/an*^ mice. In fibroblasts, Cdk5rap2 loss induces p53 Ser15 phosphorylation that correlates with decreased level of the p53 phosphatase, WIP1. Ectopic WIP1 expression reverses senescence in Cdk5rap2-depleted cells, linking senescence to WIP1 downregulation. Cdk5rap2 interacts with GSK3β, increasing inhibitory Ser9 phosphorylation in GSK3β, which phosphorylates and tags β-catenin for degradation. Thus, Cdk5rap2 loss decreases GSK3β Ser9 phosphorylation and increases GSK3β activity, reducing β-catenin that affects expression of NF-κB target genes, including WIP1. Consequently, Cdk5rap2 or β-catenin depletion downregulates WIP1. GSK3β Inhibition in Cdk5rap2-depleted cells restores β-catenin and WIP1 levels, reducing p53 Ser15 phosphorylation and preventing senescence. Conversely, WIP1 inhibition increases p53 Ser15 phosphorylation and senescence in Cdk5rap2-depleted cells lacking GSK3β activity. Senescence through GSK3β/β-catenin downregulation of WIP1 may contribute to the developmental disorders associated with Cdk5rap2 loss-of-function.

## Introduction

Cyclin-dependent kinase 5 (Cdk5) regulatory subunit associated protein 2 (Cdk5rap2) was originally identified based on its ability to interact with the Cdk5 regulatory subunit, p35^1^. Cdk5rap2 localizes to the centrosome in a dynein-dependent manner^2^ and regulates centriole engagement and centrosome cohesion during the cell cycle^3-5^. Cdk5rap2 moved into the spotlight as loss-of-function mutations in Cdk5rap2 cause primary microcephaly (MCPH), an autosomal recessive neurodevelopmental disorder characterized by small brain and cognitive deficit^6, 7^. In fact, Cdk5rap2 is most abundant at the luminal surface of the brain’s ventricular zone, particularly in cells lining the ventricular wall where neural stem and progenitor cells reside. However, since Cdk5rap2 is also expressed in other tissues, it is not surprising that loss-of-function mutations in Cdk5rap2 are further associated with other developmental disorders such as cochlear and retinal developmental defects^8^, primordial dwarfism^9, 10^, as well as Seckel syndrome, a heterogeneous, autosomal recessive disorder characterized by prenatal proportionate short stature, severe microcephaly, intellectual disability, and characteristic ‘bird-headed’ facial features^11^. However, the molecular mechanisms by which Cdk5rap2 loss-of-function mutations cause these developmental disorders remain elusive.

Cellular senescence is a state of stable cell cycle arrest in which cells remain viable and metabolically active^12^. In addition to cell cycle arrest, cell senescence is characterized by increased senescence-associated β-galactosidase (SA-β-gal) activity^13^, formation of senescence-associated heterochromatin foci (SAHF)^14^ and morphological transformation^15^. Senescence is triggered by a number of cellular stresses^16-19^. For example, activated oncogenes such as H-RAS^*G12V*^ and B-RAF^*V600E*^ induce senescence by evoking sustained anti-proliferative response^20^, a mechanism that acts as an initial barrier in preventing normal cell transformation into malignant cells. H-RAS^*G12V*^-induced senescence is associated with accumulation of DNA damage foci and activation of the p53 kinases, ataxia telangiectasia mutated (ATM), checkpoint kinase 1 (Chk1) and/or checkpoint kinase 2 (Chk2), suggesting that aberrant oncogene activation induces a DNA damage response (DDR)^20, 21^. Cell senescence can also be induced by telomere attrition and oxidative stress among others. The molecular mechanisms of senescence have been studied extensively using a variety of cell types. Phosphorylation of the tumor suppressor protein, p53, at Ser15 by p53 kinases such as ATM, Chk1 and/or Chk2 is one of the key events in p53-associated cell senescence^22, 23^. Apparently, p53 Ser15 phosphorylation stabilizes p53 by inhibiting its interaction with Mdm2, an E3 ubiquitin ligase that catalyzes polyubiquitination and subsequently induces proteasome degradation of p53^24^. However, p53 Ser15 phosphorylation is also required for p53 activation^25, 26^, which blocks cell cycle progression by inducing p21^*CIP1*^ expression^27^. Aside from phosphorylation by kinases, p53 Ser15 phosphorylation is also regulated by p53 phosphatases such as wild-type p53-induced phosphatase 1 (WIP1). In fact, WIP1 has been associated with p53-mediated cell senescence^28, 29^. For example, hematopoietic stem cells (HSC) from WIP1^-/-^ mice exhibit senescent phenotypes, impairing repopulating activity^29^. Mouse embryonic fibroblasts (MEFs)^29^ and mesenchymal stem cells^30^ from WIP1^-/-^ mice also undergo premature senescence. In primary chondrocytes, reduced WIP1 is associated with senescent phenotype, which is reversed by ectopic WIP1 expression^31^. *WIP1*, which contains an NF-κB binding site in its promoter region, is a downstream gene target of NF-κB^32^. Interestingly, β-catenin associates with NF-κB and induces the expression of NF-κB target genes^33^ such as WIP1.

Thus, our investigations examine the possibility that WIP1 expression is regulated by β-catenin and whether WIP1-associated p53-mediated senescence is linked to phenotypes related to Cdk5rap2 loss-of-function. Since β-catenin is phosphorylated by GSK3β that earmarks β-catenin for ubiquitination-mediated degradation by the proteasome pathway^34^, we further examined whether GSK3β controls our presumed β-catenin-mediated WIP1 expression. To explore these possibilities, we used two model systems: (i) the Cdk5rap2^*an/an*^ mouse model^35^, which carries exon 4 inversion in *Cdk5rap2*, and exhibits primary microcephaly that is comparable to that of the human disease, displaying proliferation and survival defects in neuronal progenitor cells^35^. Cdk5rap2^*an/an*^ homozygotes are spontaneously aborted, appearing to exhibit late embryonic lethality. (ii) BJ-5ta normal human diploid foreskin fibroblast cells, which were immortalized through forced expression of the hTERT component of telomerase. BJ-5ta cells maintain many characteristics of normal primary cells such as karyotype, growth rate, morphology, tissue-specific markers and contact inhibition. Using the Cdk5rap2^*an/an*^ mouse model and BJ-5ta cells, we demonstrate that loss of Cdk5rap2 triggers premature cell senescence. Proliferation defect and senescent phenotypes in Cdk5rap2-depleted BJ-5ta cells are recapitulated in Cdk5rap2^*an/an*^ as reduced embryonic body weights and reduced MEF growth rate, and increased SA-β-gal staining and reduced PCNA but increased p21^*CIP1*^ levels in MEFs, respectively. We propose that premature cell senescence due to Cdk5rap2 loss-of-function occurs via elevation of GSK3β activity that causes β-catenin-mediated downregulation of WIP1 and subsequent upregulation of p53 Ser15 phosphorylation.

## Results

### Loss of Cdk5rap2 induces cellular senescence

To investigate how Cdk5rap2 loss-of-function may cause proliferation defects associated with developmental disorders, we examined the effect of Cdk5rap2 depletion in BJ-5ta human fibroblasts by siRNA. As shown in Figure 1, knocking down Cdk5rap2 using two different siRNAs (#1 and #2) triggers the formation of senescence-associated heterochromatin foci (SAHF) as demonstrated by staining with DAPI (Figure 1B) and SAHF colocalization with heterochromatin protein 1γ (HP1γ, Figure 1C), a SAHF marker^36^. Senescence induced by activated H-RAS^*G12V*^ oncogene^37^ was used as positive control to detect SAHF and HP1γ staining. To support the suggestion that Cdk5rap2 depletion causes premature cellular senescence, cells transfected with Cdk5rap2 siRNA #2 (Figure 2) were monitored for the appearance of SAHF-positive cells and the expression of the senescence-associated biomarkers^36^, p21^*CIP1*^ and p16^*INK4a*^, over 5 days post transfection. As shown in Figure 2A, the number of SAHF positive cells increases markedly in the first 3 days after transfection with Cdk5rap2 siRNA #2, reaching peak levels on days 3-5, a period when no or only a modest number of SAHF positive cells was observed in those transfected with control siRNA. The increase in number of SAHF positive cells upon Cdk5rap2 depletion coincides with increased expression of p21^*CIP1*^ and p16^*INK4a*^ (Figure 2B). Together with the noticeable staining for SA-β-gal, another marker of cellular senescence^38^, 3 days post-transfection with Cdk5rap2 siRNA #2 (Figure 2C), our findings indicate that loss of Cdk5rap2 induces premature cellular senescence.

**Figure 1.**
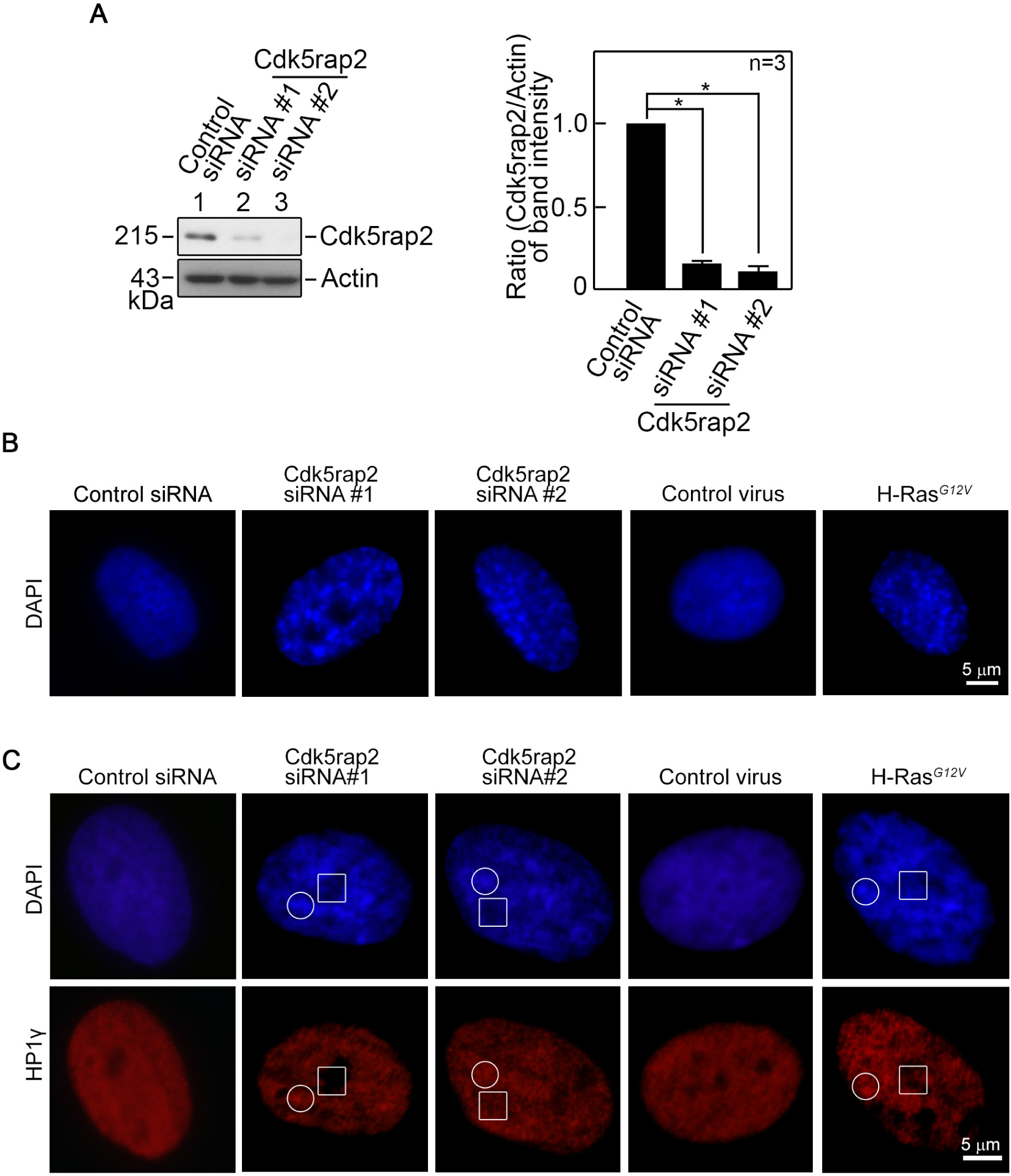
Cdk5rap2 loss triggers SAHF formation. A. Depletion of Cdk5rap2 in BJ-5ta human diploid foreskin fibroblasts. Lysates of cells transfected with Cdk5rap2 siRNA for 3 days were analyzed by SDS-PAGE and immunoblotting for Cdk5rap2 (left panel). Actin blot was used as loading control. Representative blots are from one of three independent experiments (n=3) showing similar results. Ratios of levels of Cdk5rap2 vs actin (right panel) were calculated following densitometric analysis of blots using NIH Image J 1.61. Standard deviation for the 3 independent sets of experiments was calculated based on the ratios of the densitometric levels of Cdk5rap2 vs actin, with values from cells transfected with control siRNA normalized to 1.0. *p<0.001. B and C. Cdk5rap2 loss causes formation of SAHF. BJ-5ta cells transfected with Cdk5rap2 siRNA for 3 days were stained with DAPI (B) and HP1γ antibody (C), and subjected to microscopic examination. SAHF formation induced by infecting adenovirus carrying H-RAS^*G12V*^ into BJ cells was used as positive control. Representative images of control and Cdk5rap2-depleted cells, and cells infected with adenovirus carrying GFP alone or H-RAS^*G12V*^ are shown with DAPI and HP1γ stain-enriched and -devoid regions highlighted with open circles and squares, respectively.

**Figure 2.**
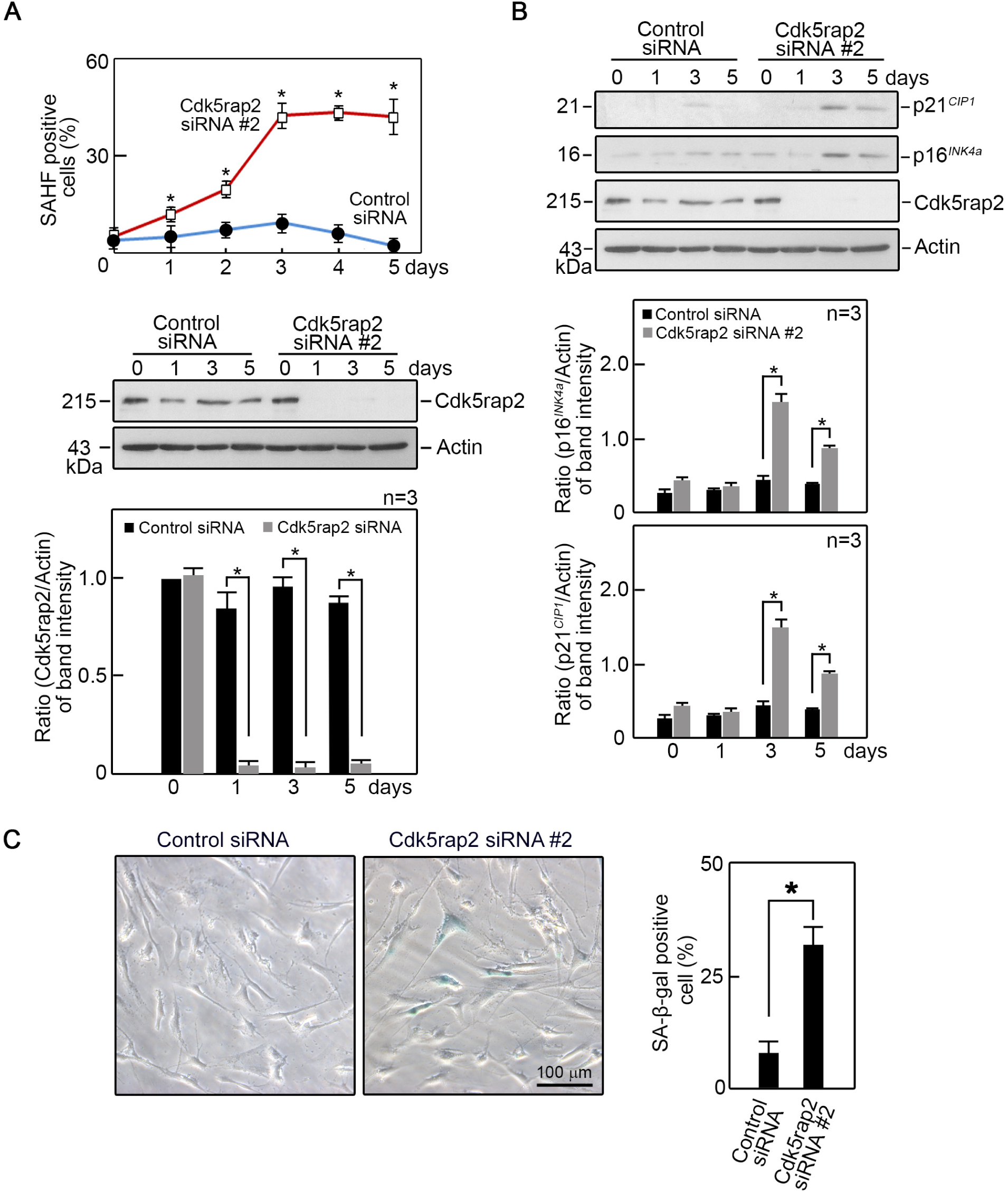
Increased number of SAHF positive Cdk5rap2-depleted cells coincides with expression of p21^*CIP1*^ and p16^*INK4a*^. BJ-5ta cells transfected with Cdk5rap2 siRNA #2 for 5 days were analyzed (i) for SAHF positive cells (A, upper panel) and by SDS-PAGE and immunoblotting for (ii) Cdk5rap2 and actin (A, middle panel), and (iii) p21^*CIP1*^ and p16^*INK4a*^ as well as Cdk5rap2 and actin (B, upper panel). The number of SAHF positive cells was assessed in ∼200 cells per treatment group in each of 3 independent experiments (n=3). *p<0.01. Actin was used as loading control for immunoblotting. Representative blots are from one of three independent experiments (n=3) showing similar results. Ratios of levels of Cdk5rap2, p21^*CIP1*^ and p16^*INK4a*^ vs actin and standard deviation for the 3 independent sets of experiments (A, lower panel and B, middle and lower panels) were calculated as described for Cdk5rap2 vs actin in Figure 1, with values from cells transfected with control siRNA at day 0 normalized to 1.0. *p<0.02. C. Cells transfected with Cdk5rap2 siRNA #2 or control siRNA were subjected to SA-β-gal staining 3 days post-transfection. Representative images (left panel) are from one of three independent experiments (n=3) showing similar staining patterns. The number of SA-β-gal positive cells was assessed in ∼200 cells per treatment group in each of the 3 independent experiments (n=3). *p=0.0005.

Increased levels of SAHF as well as p21^*CIP1*^ and p16^*INK4a*^, which are also molecular inhibitors of cell cycle Cdks, upon loss of Cdk5rap2 led us to test whether loss of Cdk5rap2 affects cell proliferation. Indeed, we found that cells transfected with Cdk5rap2 siRNA #2 proliferate at a slower rate compared to cells transfected with control siRNA (Figure 3A). Flow cytometry analysis show greater population of cells at G0/G1 but reduced population of cells at S in those transfected with Cdk5rap2 siRNA #2 compared to those transfected with control siRNA (Figure 3B). In addition, reduced number of Ki-67 positive cells were observed in those transfected with Cdk5rap2 siRNA #2 compared to those transfected with control siRNA (Figure 3C). Altogether, these findings indicate that Cdk5rap2 loss triggers senescence-associated proliferation arrest.

**Figure 3.**
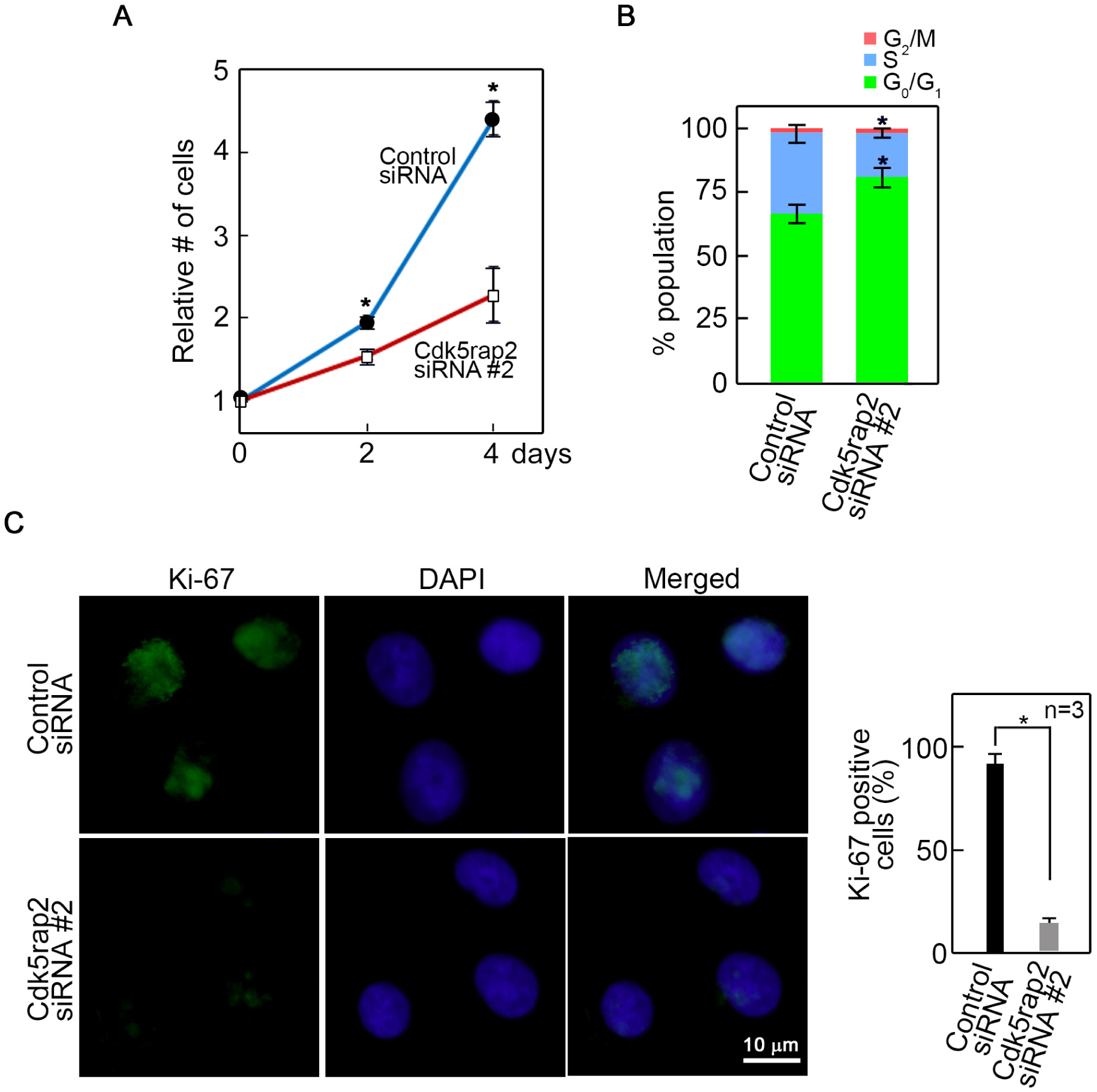
Cdk5rap2 loss causes decreased cell proliferation. BJ-5ta cells transfected with Cdk5rap2 siRNA #2 were subjected to (A) cell viability assay, (B) cell cycle distribution analysis by flow cytometry, and (C) measurement of Ki-67 positive cells as described in Materials and Methods. Data represent means ± SD from three separate experiments (n=3). *p<0.01 (in A and B). The percentage (%) of Ki-67 positive cells (C, right panel) were assessed in ∼200 cells per treatment group in each of the 3 independent experiments (n=3). *p=0.001.

### Mouse embryonic fibroblast cells (MEFs) isolated from Cdk5rap2^*an/an*^ mice, which have reduced body weights, exhibit senescence-associated phenotypes

The availability of Hertwig’s anemia (an) mutant mice (Cdk5rap2^*an/an*^) that carry exon 4 inversion in the *Cdk5rap2* gene, allowed us to test whether such Cdk5rap2^*an/an*^ mutation affects body weight and whether the senescent-associated phenotypes observed in cells lacking Cdk5rap2 are also exhibited by Cdk5rap2^*an/an*^ MEFs. Since Cdk5rap2^*an/an*^ mice have high incidence of death in late embryonic stages, we initially isolated embryonic day 12.5 (E12.5) Cdk5rap2^*+/+*^, Cdk5rap2^*+/an*^ and Cdk5rap2^*an/an*^ embryos from pregnant Cdk5rap2^*+/an*^ mice and body weights were compared. As shown in Figure 4A, within the same litter, the Cdk5rap2^*an/an*^ embryo weighs less than the Cdk5rap2^*+/+*^ and Cdk5rap2^*+/an*^ embryos. E17.5 embryos from different litters also show lower average weights of Cdk5rap2^*an/an*^ compared to Cdk5rap2^*+/+*^ and Cdk5rap2^*+/an*^ embryos (Figure 4B, left panel). Similar observations were further noted for average weights of E12.5 to E17.5 Cdk5rap2^*+/+*^, Cdk5rap2^*+/an*^ and Cdk5rap2^*an/an*^ embryos (Figure 4B, right panel). Next, MEFs isolated from E12.5 Cdk5rap2^*+/+*^, Cdk5rap2^*+/an*^ and Cdk5rap2^*an/an*^ embryos were examined for the appearance of SA-β-gal positive cells (Figure 4C, left panel). Analysis of SA-β-gal staining show that the number of SA-β-gal positive cells in Cdk5rap2^*an/an*^ MEFs is remarkably greater than those in Cdk5rap2^*+/+*^ and Cdk5rap2^*+/an*^ MEFs (Figure 4C, right panel). In addition, Cdk5rap2^*an/an*^ MEFs exhibit decreased proliferation (Figure 4D), which coincides with reduced PCNA but increased p21^*CIP1*^ levels (Figure 4E) compared to Cdk5rap2^*+/+*^ and Cdk5rap2^*+/an*^ MEFs. Thus, the senescence-associated phenotypes that we observed in Cdk5rap2-depleted BJ-5ta cells is recapitulated in *ex vivo* Cdk5rap2^*an/an*^ MEFs.

**Figure 4.**
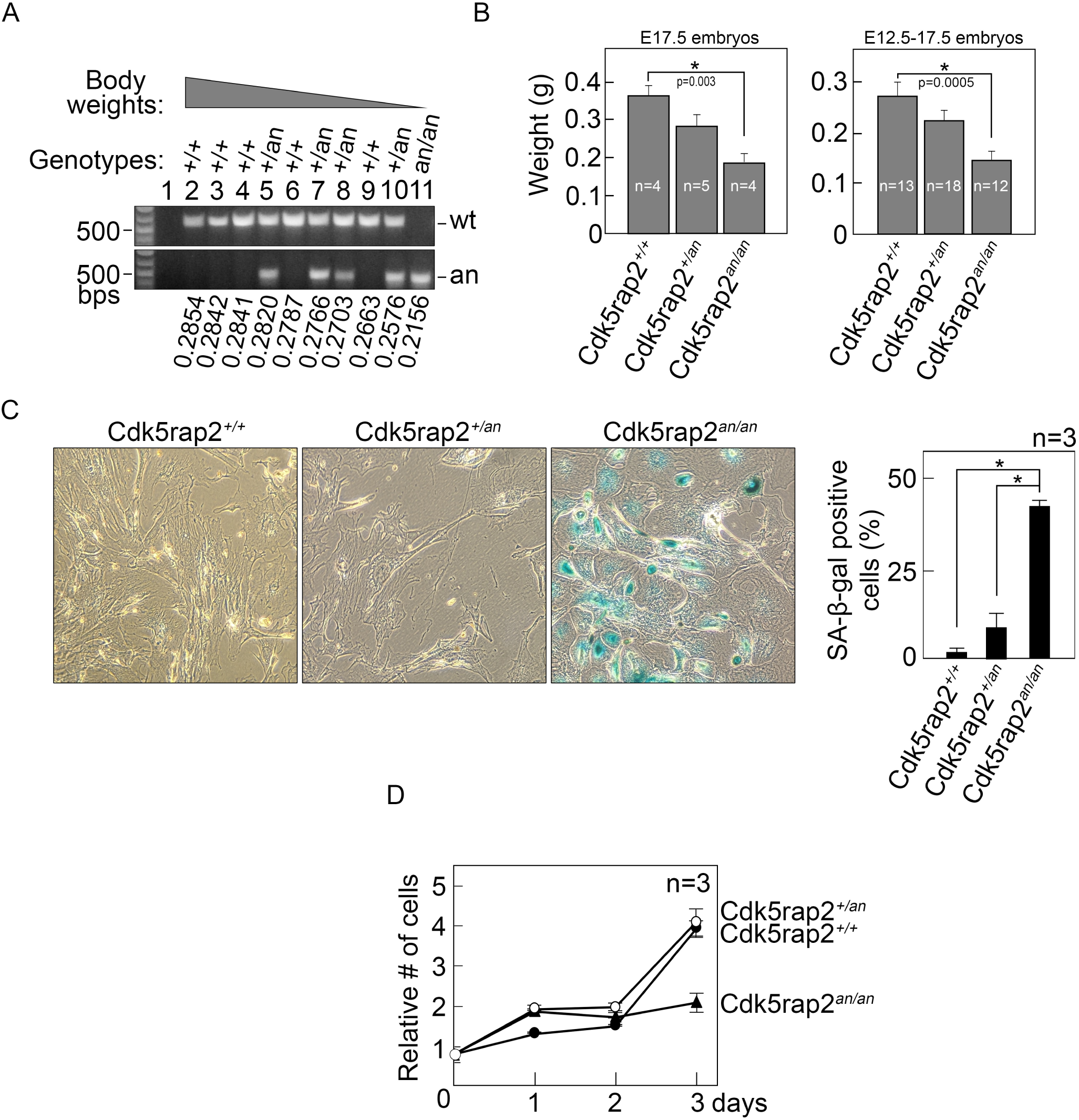
Cdk5rap2^*an/an*^ embryos have reduced body weight and Cdk5rap2^*an/an*^ MEFs show senescent phenotype. A. Genomic DNAs from E12.5 Cdk5rap2^*+/+*^, Cdk5rap2^*+/an*^ and Cdk5rap2^*an/an*^ littermates obtained from a Cdk5rap2^*+/an*^ pregnant mouse were subjected to genotyping by PCR as described in Materials and Methods. Lane 1 is a negative control where no PCR template was added. Embryos were numbered (2-11) according to their weights (heaviest to lightest). B. Average weights of isolated E17.5 (left panel) and E12.5 to E17.5 (right panel) Cdk5rap2^*+/+*^, Cdk5rap2^*+/an*^ and Cdk5rap2^*an/an*^ embryos from different litters are shown. C. MEFs isolated from Cdk5rap2^*+/+*^, Cdk5rap2^*+/an*^ and Cdk5rap2^*an/an*^ embryos were subjected to SA-β-gal staining. Representative images (left panel) are from one of three independent experiments (n=3) showing similar staining patterns. The number of SA-β-gal positive cells was assessed in ∼200 cells per treatment group in each of the 3 independent experiments (n=3). *p=0.0002. D. Growth of MEFs obtained from Cdk5rap2^*+/+*^, Cdk5rap2^*+/an*^ and Cdk5rap2^*an/an*^ embryos was analyzed by cell viability assay. E. Lysates of MEFs isolated from Cdk5rap2^*+/+*^, Cdk5rap2^*+/an*^ and Cdk5rap2^*an/an*^ embryos were analyzed by SDS-PAGE and immunoblotting for Cdk5rap2, PCNA and p21^*CIP1*^. TCE staining was used to assess protein loading.

### Cell senescence due to loss of Cdk5rap2 is linked to elevated p53 Ser15 phosphorylation via WIP1 downregulation

p53 plays a key role in triggering the expression of SA-β-gal. In addition, upon DNA damage and other types of stress that induce cellular senescence^25, 39^, p53 is phosphorylated at Ser15 by protein kinases such as ATM, Chk1, and Chk2^22-24^. Thus, we examined the status of p53 Ser15 phosphorylation in BJ-5ta cells depleted of Cdk5rap2. By immunoblotting using a phosphoSer15 p53 antibody, we observed that loss of Cdk5rap2 causes noticeable increase in p53 Ser15 phosphorylation, which coincides with elevation of p21^*CIP1*^ level (Figure 5A). Next, we tested whether senescence due to Cdk5rap2 loss involves ATM, Chk1 and/or Chk2 activation. Phosphorylation of ATM Ser1981, Chk1 Ser345 and Chk2 Thr68 was examined in Cdk5rap2-depleted cells by immunoblotting. Cells infected with H-RAS^*G12V*^ oncogene^37^ was used as positive control to detect senescence-associated activation of ATM, Chk1 and Chk2. Interestingly, we did not detect activation of ATM, Chk1 and Chk2 in Cdk5rap2-depleted cells (Figure 5B). Unexpectedly, we observed reduced level of Cdk5rap2 in cells infected with activated H-RAS^*G12V*^. The mechanism by which this happens requires further investigation but is beyond the scope of the current study. Since phosphorylation of p53 Ser15 is also regulated by protein phosphatases such as WIP1, a critical senescence regulator^28, 29^ that has been associated with p53 Ser15 dephosphorylation, we examined the possibility that loss of Cdk5rap2 affects WIP1 expression. By qRT-PCR and immunoblotting, we found that WIP1 phosphatase is downregulated at both the mRNA (Figure 5C, left panel) and protein (Figure 5C, right upper and lower panels) levels in cells depleted of Cdk5rap2. Taken together, our findings indicate that premature cellular senescence due to loss of Cdk5rap2 may be induced by p53 phosphorylation at Ser15 via downregulation of WIP1, and that p53-mediated senescence due to Cdk5rap2 loss is independent of ATM, Chk1 and Chk2 activation.

**Figure 5.**
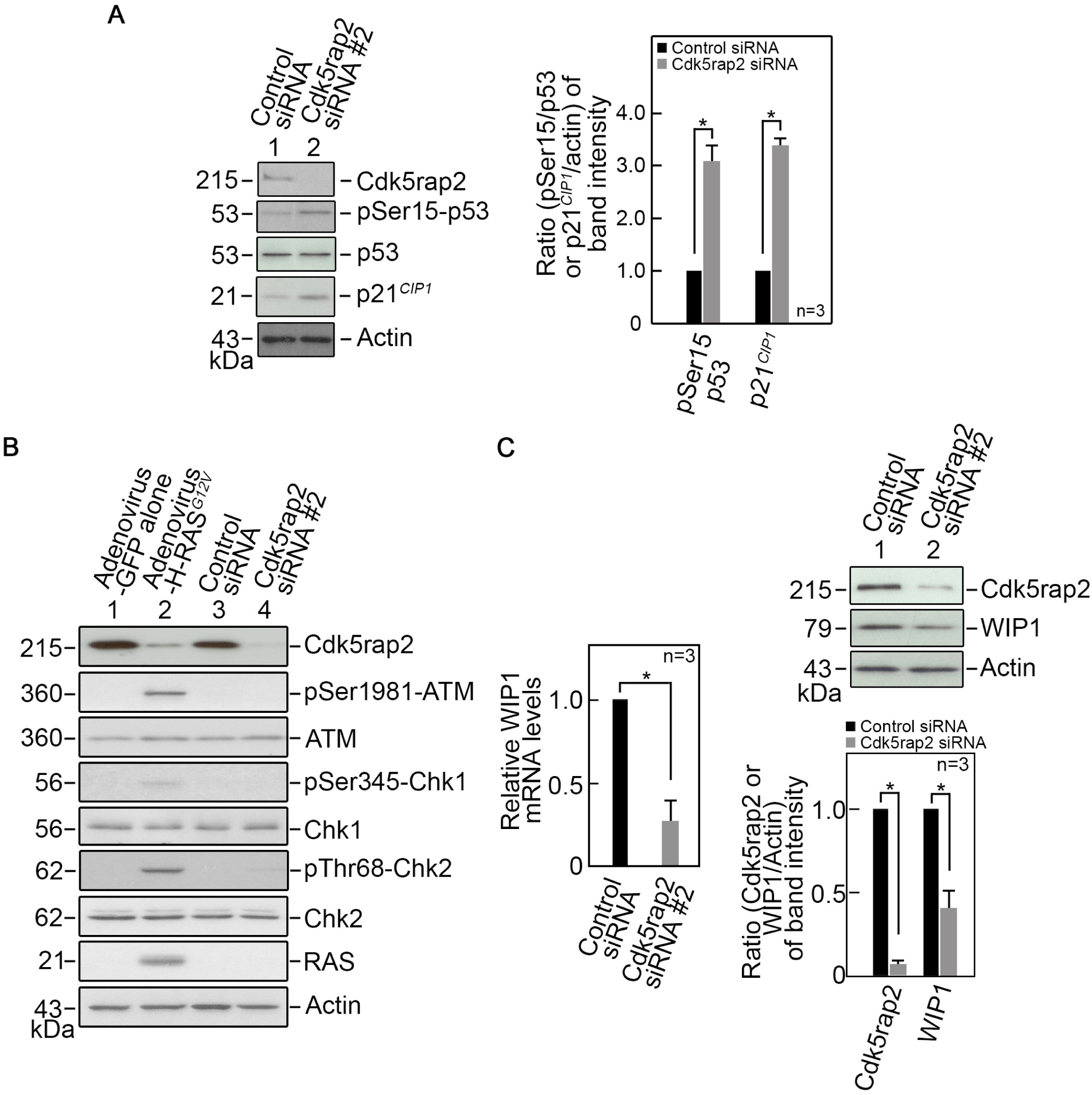
Cdk5rap2 loss triggers cellular senescence through downregulation of WIP1-mediated p53 activation. A. BJ-5ta cells depleted of Cdk5rap2 show increased p53 phosphorylation at Ser15, which coincides with increased level of p21^*CIP1*^. Lysates of cells transfected with Cdk5rap2 siRNA #2 for 3 days were analyzed by SDS-PAGE and immunoblotting for Cdk5rap2, phosphoSer15 p53 (pSer15-p53), p53, p21^*CIP1*^ and actin (A, left panel). Actin blot was used as loading control. Representative blots are from one of three independent experiments (n=3) showing similar results. Ratios of levels of phosphoSer15 p53 (pSer15-p53) and p21^*CIP1*^ vs actin and standard deviation for the 3 independent sets of experiments (A, right panel) were calculated as described for Cdk5rap2 vs actin in Figure 1, with values from cells transfected with control siRNA normalized to 1.0. *p<0.005. B. Lysates of cells infected with adenovirus carrying H-RAS^*G12V*^ or transfected with Cdk5rap2 siRNA #2 for 3 days were analyzed by SDS-PAGE and immunoblotting for Cdk5rap2, phosphoSer1982-ATM (pSer1982-ATM), ATM, phosphoSer345-Chk1 (pSer345-Chk1), Chk1, phosphoThr68-Chk2 (pThr68-Chk2), Chk2, H-RAS and Actin. Actin blot was used as loading control. C. Cdk5rap2 loss represses WIP1 mRNA and protein expression. Levels of WIP1 mRNA in cells transfected with Cdk5rap2 siRNA #2 were determined by qRT-PCR using isolated total RNA as template. GAPDH was used for normalization. Relative levels of WIP1 mRNA (left panel) were calculated using the 2^−ΔΔCT^ method. Relative levels from cells transfected with control siRNA were normalized to 1.0. Data represent means ± SD of calculated ratios from three independent experiments (n=3). *p=0.002. WIP1 protein expression was analyzed by immunoblotting (upper right panel). Actin blot was used as loading control. Representative blots from one of three independent experiments (n=3) showing similar results are shown. Ratios of levels of WIP1 vs actin and standard deviation for the 3 independent sets of experiments (lower right panel) were calculated as described for Cdk5rap2 vs actin in Figure 1, with values from cells transfected with control siRNA normalized to 1.0. *p<0.01.

While we note that p53-associated cell senescence induced by loss of Cdk5rap2 is distinct from the p53-mediated senescence induced by activated H-RAS^*G12V*^, which is often associated with activation of the DNA damage response (DDR)^19, 21-23^, we sought to examine the possibility that senescence induced by Cdk5rap2 loss involves DDR. To do so, we performed immunofluorescence microscopy to test whether loss of Cdk5rap2 in BJ-5ta cells triggers the formation of γH2AX foci, which was assessed by staining with histone H2AX phosphoSer139 antibody (supplementary Figure 1). Cells infected with activated H-RAS^*G12V*^ and cells treated with 4Gy ionizing radiation (IR), which show 96.7% and 100%, respectively, of cells positive for γH2AX foci served as positive controls. Cells infected with an empty vector and cells transfected with control siRNA served as negative controls. Examination of Cdk5rap2-depleted cells revealed no γH2AX foci, indicating that p53-associated senescence that is linked to WIP1 downregulation in Cdk5rap2-depleted cells does not involve DDR activation.

To examine the idea that WIP1 loss in Cdk5rap2-depleted cells accounts for p53 Ser15 phosphorylation that leads to cellular senescence, BJ-5ta cells depleted of WIP1 by siRNA were analyzed by SDS-PAGE and immunoblotting for WIP1, p53 phophoSer15, total p53 and p21^*CIP1*^ as well as SA-β-gal staining. Consistent with our observations in Cdk5rap2-depleted cells, WIP1-depleted cells show p53 Ser15 phosphorylation and elevation of p21^*CIP1*^ level (Figure 6A), and increased number of SA-β-gal positive cells (Figure 6B). To establish a link between senescence induced by Cdk5rap2 loss and downregulated WIP1 expression, BJ-5ta cells were co-transfected with Cdk5rap2 siRNA #2 and pReceiver-M02 vector carrying WIP1. As shown in Figure 7, ectopic WIP1 overexpression in Cdk5rap2-depleted cells (Figure 7A) reverses the appearance of SA-β-gal positive cells induced by Cdk5rap2 loss (Figure 7B), supporting our view that senescence due to loss of Cdk5rap2 is linked to downregulated WIP1 expression.

**Figure 6.**
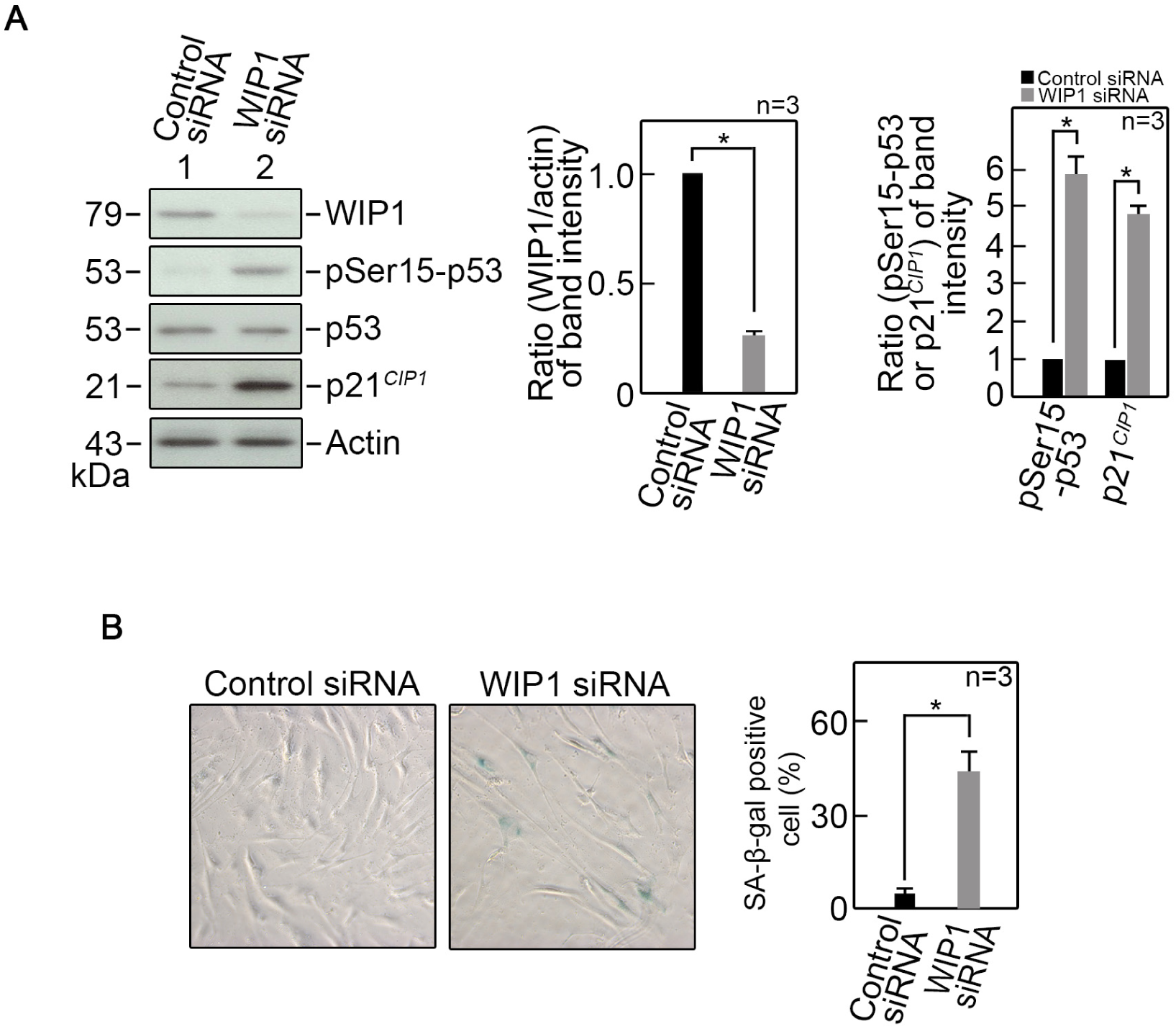
WIP1 loss triggers cellular senescence through p53 activation. BJ-5ta cells transfected with WIP1 siRNA were analyzed by (A) SDS-PAGE and immunoblotting for WIP1, phophoSer15-p53, p53, and p21^*CIP1*^ (left panel), and (B) SA-β-gal staining (left panel). Actin was used as loading control for immunoblotting. Representative blots are from one of three independent experiments (n=3) showing similar results. Ratios of levels of WIP1 vs actin (A, middle panel) and phophoSer15-p53 vs p53 and p21^*CIP1*^ vs actin (A, right panel) and standard deviation for the 3 independent sets of experiments were calculated as described for Cdk5rap2 vs actin in Figure 1, with values from cells transfected with control siRNA normalized to 1.0. *p<0.005. Representative images of SA-β-gal staining in B (left panel) are from one of three independent experiments (n=3) showing similar staining patterns. The number of SA-β-gal positive cells (B, right panel) was assessed in ∼200 cells per treatment group in each of the 3 independent experiments (n=3). *p=0.00016.

**Figure 7.**
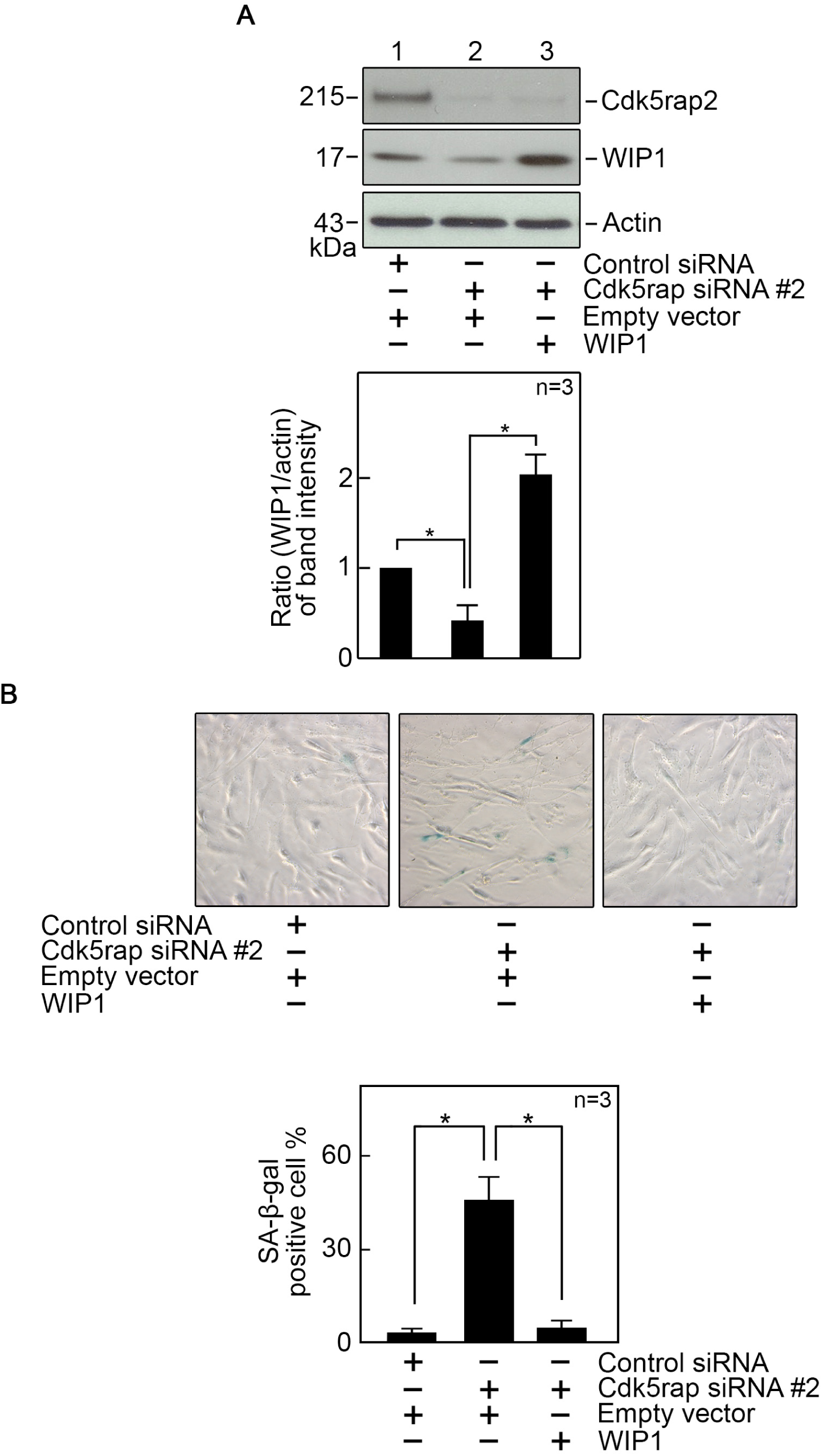
Ectopic expression of WIP1 reverses the senescent phenotypes observed in cells depleted of Cdk5rap2. Lysates of BJ-5ta cells co-transfected with Cdk5rap2 siRNA #2 and pReceiver-M02 carrying WIP1 were analyzed by SDS-PAGE and immunoblotting for Cdk5rap2 and WIP1 (A, upper panel), and SA-β-gal staining (B, upper panel) 72 hrs post-transfection. Actin was used as loading control for immunoblotting. Representative blots are from one of three independent experiments (n=3) showing similar results. Ratios of levels of WIP1 vs actin and standard deviation for the 3 independent sets of experiments (A, lower panel) were calculated as described for Cdk5rap2 vs actin in Figure 1, with values from cells co-transfected with control siRNA and empty vector normalized to 1.0. *p<0.05. Representative images of SA-β-gal staining in B (upper panel) are from one of three independent experiments (n=3) showing similar staining patterns. The number of SA-β-gal positive cells (B, lower panel) was assessed in ∼100 cells per treatment group in each of the 3 independent experiments (n=3). *p<0.001.

### Cdk5rap2 regulates WIP1 expression through β-catenin

As indicated above, we found that loss of Cdk5rap2 causes downregulation of WIP1 expression at both the mRNA and protein levels. The WIP1 promoter contains a nuclear factor-κB (NF-κB) binding site^32^ (Figure 8A) through which NF-κB positively regulates WIP1 expression^32^, and β-catenin associates with NF-κB, affecting the expression of NF-κB target genes^33^. These apparent interactions led us to test whether loss of Cdk5rap2, which causes reduced WIP1 level, influences β-catenin expression. As shown in Figure 8B, loss of Cdk5rap2 reduces β-catenin level in the nucleus where β-catenin associates with NF-κB. Depletion of β-catenin by siRNA downregulates WIP1 level (Figure 8C), suggesting that Cdk5rap2 loss may reduce WIP1 level through downregulation of β-catenin.

**Figure 8.**
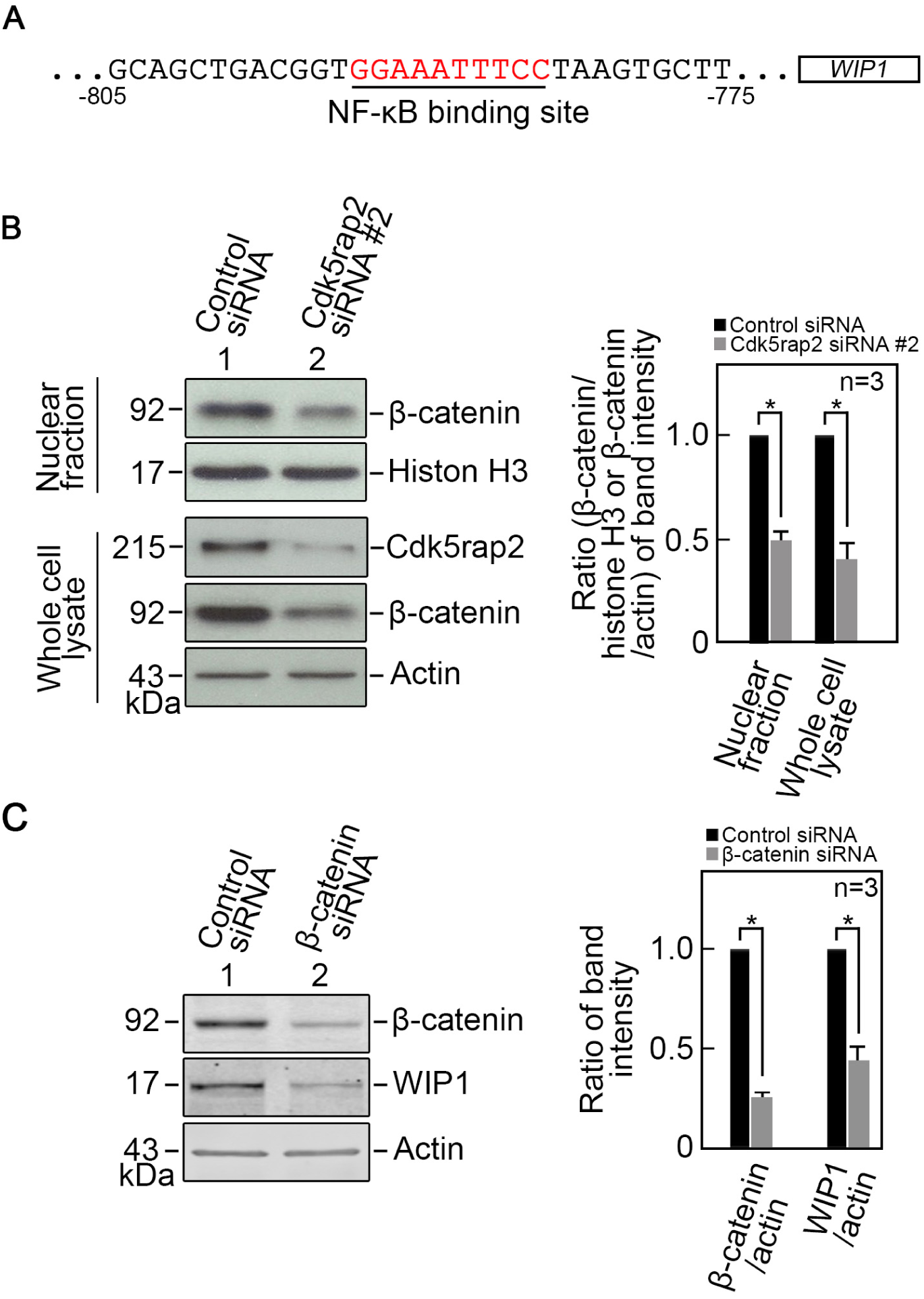
Cdk5rap2 loss reduces β-catenin level. A. The WIP1 promoter contains an NF-κB binding site^32^. BJ-5ta cells transfected with Cdk5rap2 siRNA #2 (B), or β-catenin siRNA (C) for 3 days were analyzed by SDS-PAGE and immunoblotting. Nuclear fractions and whole cell lysates in (B) were probed for β-catenin and histone H3 (nuclear fraction) or Cdk5rap2 and actin (whole cell lysate), and whole cell lysates in (C) were probed for β-catenin, WIP1 and actin. Actin blot was used as loading control. Representative blots are from one of three independent experiments (n=3) showing similar results. Ratios of levels of β-catenin vs histone H3 (in nuclear fraction) or actin (in whole cell lysate) in (B) and β-catenin or WIP1 vs actin in (C), right panels, and standard deviation for the 3 independent sets of experiments were calculated as described for Cdk5rap2 vs actin in Figure 1, with values from cells transfected with control siRNA normalized to 1.0. *p<0.05 (in B). *p<0.005 (in C).

### Cdk5rap2 interacts with glycogen synthase kinase-3β (GSK3β) and such interaction inhibits GSK3β activity

Because GSK3β phosphorylates β-catenin, targeting β-catenin for ubiquitination-mediated degradation by the proteasome pathway^34^, we examined whether Cdk5rap2 interacts with GSK3β and affects GSK3β activity. To do so, lysates of BJ-5ta cells co-transfected with Myc-tagged Cdk5rap2 and FLAG-tagged GSK3β were subjected to immunoprecipitation using FLAG or Myc antibody then analyzed for Cdk5rap2 and GSK3β co-immunoprecipitation by immunoblotting both immunoprecipitates with Myc and FLAG antibodies. Fig. 9A shows reciprocal co-immunoprecipitation of Cdk5rap2 and GSK3β. To determine whether Cdk5rap2 interacts directly with GSK3β, GST-Cdk5rap2 purified from baculovirus-infected Sf9 insect cells (supplementary Figure 2) was bound to GSH-agarose beads then mixed with purified GSK3β. Analysis of the GST pulled-down complex by SDS-PAGE and immunoblotting using GST or GSK3β antibody show the presence of both GST-Cdk5rap2 and GSK3β (Figure 9B), indicating direct interaction between these proteins. GSH-agarose-bound GST mixed with GSK3β was used as negative control. We then sought to determine whether Cdk5rap2 interaction with GSK3β affects GSK3β activity. Since GSK3β phosphorylation at Ser9 inhibits its activity^34, 40^, we first examined whether Cdk5rap2 loss influences the state of GSK3β Ser9 phosphorylation. As shown in Figure 9C, cells transfected with Cdk5rap2 siRNA #2 have reduced GSK3β Ser9 phosphorylation compared to cells transfected with control siRNA, suggesting that Cdk5rap2 interaction with GSK3β could affect GSK3β activity. Indeed, by *in vitro* GSK3β kinase assay, we found that the GSK3β immunoprecipitate from cells transfected with Cdk5rap2 siRNA #2 show greater GSK3β activity compared to cells transfected with control siRNA (Figure 9D). To examine a link between increased GSK3β activity and reduced β-catenin and WIP1 levels in Cdk5rap2-depleted cells, we tested whether inhibition of GSK3β with TWS119 restores β-catenin and WIP1 levels in these cells. Western blot analysis shows that TWS119 restores β-catenin and WIP1 levels in cells transfected with Cdk5rap2 siRNA #2 (Figure 9E), supporting a link between increased GSK3β activity and reduced β-catenin and WIP1 levels upon loss of Cdk5rap2.

**Figure 9.**
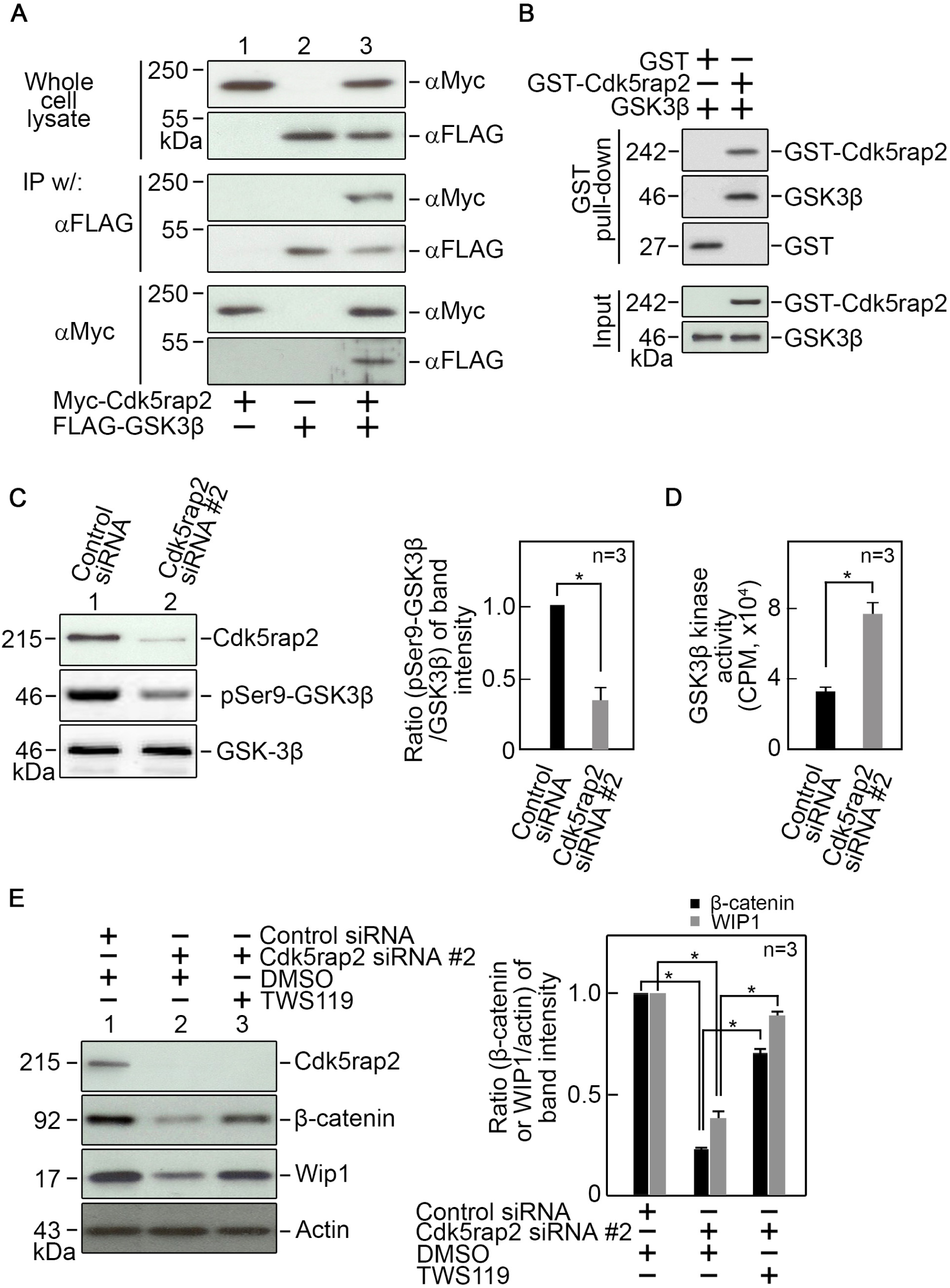
GSK3β interaction with Cdk5rap2 inhibits its kinase activity. A. Lysates of BJ-5ta cells co-transfected with Myc-tagged Cdk5rap2 and FLAG-tagged GSK3β (upper panel) were subjected to immunoprecipitation (IP) using FLAG antibody (middle panel) or Myc antibody (lower panel) then analyzed for Cdk5rap2 and GSK3β co-immunoprecipitation by immunoblotting both IPs with Myc and FLAG antibodies. B. Cdk5rap2 directly interacts with GSK3β. GST-Cdk5rap2 bound to GSH-agarose beads was mixed with GSK3β as described in Materials and Methods. The resulting complexes were washed extensively then analyzed by SDS-PAGE and immunoblotting for GST and GSK3β. C. Lysates of cells transfected with Cdk5rap2 siRNA #2 for 3 days were analyzed by SDS-PAGE and immunoblotting for Cdk5rap2, phosphoSer9-GSK3β and GSK3β (left panel). Representative blots are from one of three independent experiments (n=3) showing similar results. Ratios of levels of phosphoSer9-GSK3β vs GSK3β and standard deviation for the 3 independent sets of experiments (right panel) were calculated as described for Cdk5rap2 vs actin in Figure 1, with values from cells transfected with control siRNA normalized to 1.0. *p=0.01. D. Lysates of cells transfected with Cdk5rap2 siRNA #2 for 3 days were subjected to IP using GSK3β antibody. The IPs were then examined for *in vitro* GSK3β kinase activity as described in Materials and Methods. Data represent means ± SD from three independent experiments (n=3). *p=0.0027. E. Inhibition of GSK3β activity with TWS119 in cells depleted of Cdk5rap2 restores β-catenin and WIP1 expression (left panel). Representative blots from one of three independent experiments (n=3) showing similar results are shown. Ratios of levels of β-catenin and WIP1 vs actin and standard deviation for the 3 independent sets of experiments (right panel) were calculated as described for Cdk5rap2 vs actin in Figure 1, with values from cells transfected with control siRNA and treated with DMSO normalized to 1.0. *p<0.05.

### Cdk5rap2 loss induces senescence through upregulation of GSK3β activity and subsequent downregulation of β-catenin-mediated WIP1 expression

Our next step was to examine whether inhibition of GSK3β with TWS119, which restores β-catenin and WIP1 levels in BJ-5ta cells depleted of Cdk5rap2, inhibits senescence in these cells. We found that treatment of BJ-5ta cells with TWS119, which increases β-catenin and WIP1 levels and reduces p53 Ser15 phosphorylation in cells transfected with Cdk5rap2 siRNA #2 (Figure 10A), results in loss of SA-β-gal staining in these cells (Figure 10B), suggesting that inhibition of GSK3β prevents senescence in Cdk5rap2-depleted cells through upregulation of β-catenin and WIP1 and subsequent downregulation of p53 Ser15 phosphorylation. We then examined the effect of inhibition of WIP1 activity with GSK2830371 in Cdk5rap2-depleted cells treated with TWS119. Figure 10A shows that GSK2830371 has no effect on β-catenin and WIP1 levels in these cells but causes an increase in p53 Ser15 phosphorylation and thus, as shown in Figure 10B, an increase in SA-β-gal staining. These findings support our view that GSK3β acts upstream of β-catenin as well as WIP1 and, therefore, while GSK3β inhibition prevents senescence in Cdk5rap2-depleted cells by increasing β-catenin and WIP1 levels, reducing p53 Ser15 phosphorylation, direct inactivation of WIP1 in cells lacking Cdk5rap2 and GSK3β activity results in p53 Ser15 phosphorylation and increased senescence.

**Figure 10.**
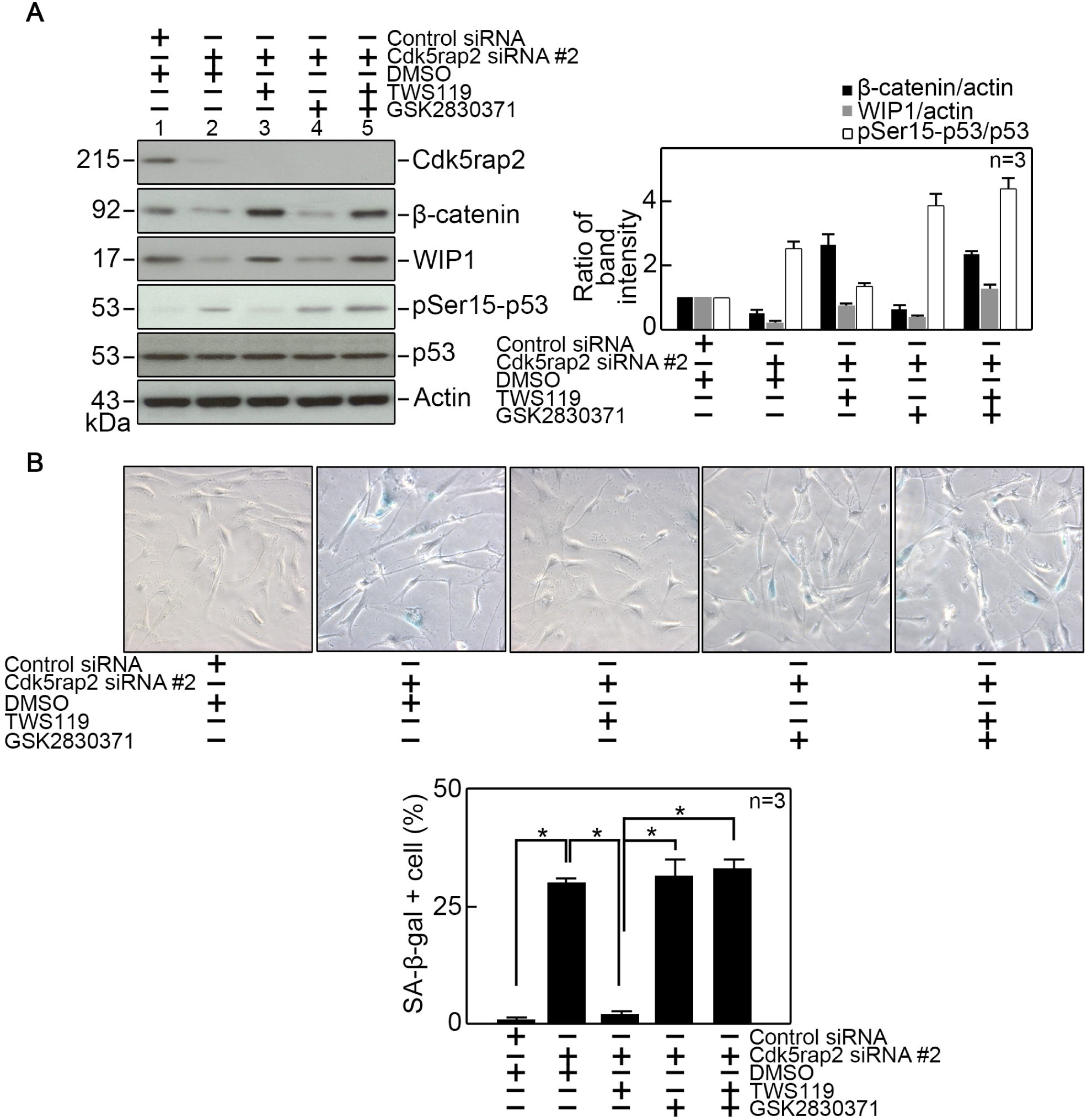
Inhibition of GSK3β in Cdk5rap2-depleted cells inhibits senescence that is induced upon inhibition of WIP1 activity. A. BJ-5ta cells transfected with Cdk5rap2 siRNA #2 were treated with TWS119 and/or GSK2830371, a potent inhibitor of WIP1 activity. Cell lysates were then analyzed by SDS-PAGE and immunoblotting for Cdk5rap2, β-catenin, WIP1, phosphoSer15-p53, p53 and actin (left panel). Actin blot was used as loading control. Representative blots from one of three independent experiments (n=3) showing similar results are shown. Ratios of levels of β-catenin and WIP1 vs actin and phosphoSer15-p53 vs p53, and standard deviation for the 3 independent sets of experiments (right panel) were calculated as described for Cdk5rap2 vs actin in Figure 1, with values from cells transfected with control siRNA and treated with DMSO normalized to 1.0. *p<0.05. B. Representative images of SA-β-gal staining (upper panel) are from one of three independent experiments (n=3) showing similar staining patterns. The number of SA-β-gal positive cells (lower panel) was assessed in ∼100 cells per treatment group in each of the 3 independent experiments (n=3). *p<0.05.

Promoter bashing analysis by transfecting a luciferase reporter vector carrying wt WIP1 promoter (pGL3-WIP1, supplementary Figure 3A) into cells transfected with Cdk5rap2 siRNA #2 (Supplementary Figure 3B) or β-catenin siRNA (Supplementary Figure 3C) show reduced luciferase activity in these cells compared to cells transfected with control siRNA (Supplementary Figure 3, B &C). Conversely, transfection with WIP1 lacking its NF-κB binding site (pGL3-WIP1-ΔκB; Supplementary Figure 3A) has no effect on luciferase activity in Cdk5rap2-or β-catenin-depleted cells. Treatment with TWS119 restored luciferase activity in Cdk5rap2-depleted cells transfected with pGL3-WIP1 (Supplementary Figure 3B, right panel), further suggesting that senescence due to Cdk5rap2 loss is controlled by GSK3β through regulation of the WIP1 promoter via β-catenin.

## Discussion

In this study, we provide evidence that Cdk5rap2 loss induces premature cell senescence through upregulation of GSK3β activity, which causes downregulation of β-catenin-modulated WIP1 level and subsequent maintenance of p53 activation through p53 Ser15 phosphorylation. These were demonstrated using BJ-5ta cells as well as Cdk5rap2^*an/an*^ mice embryos and their corresponding *ex vivo* MEF cells. In BJ-5ta cells, we eliminated the possibility of off-target effects of siRNA used to deplete Cdk5rap2 by using two different Cdk5rap2 siRNAs (#1 and #2) that both successfully depleted Cdk5rap2 and induced similar senescent phenotypes that were not observed in cells transfected with control siRNA. For subsequent experiments, Cdk5rap2 siRNA #2, which causes greater depletion of Cdk5rap2 was used. Premature BJ-5ta cell senescence due to Cdk5rap2 loss was verified by an increase in number of SAHF positive cells, upregulation of the senescence-associated genes, p21^*CIP1*^ and p16^*INK4a* 41^, and increased staining for SA-β-gal. These senescent phenotypes observed in Cdk5rap2-depleted BJ-5ta cells are recapitulated in *ex vivo* MEFs isolated from Cdk5rap2^*an/an*^ embryos, and consistent with a previous observation that MEFs isolated from mice carrying a frameshift loss-of-function mutation (R1334SfsX5)^6, 42^ in Cdk5rap2 do not grow to confluence after passages 8 to 10^43^. However, the latter finding was not further investigated. In BJ-5ta cells, proliferation defect in Cdk5rap2-depleted cells was demonstrated by increased population of cells at G0/G1 and reduced population of cells at S phase, indicating reduced cell proliferation. In Cdk5rap2^*an/an*^, cell proliferation defect is manifested by reduced body weights in E12.5, E17.5 as well as combined E12.5, E14.5, E15.5 and E17.5 embryos compared to Cdk5rap2^*+/+*^ and Cdk5rap2^*+/an*^.

Characterization of senescence upon loss of Cdk5rap2 in BJ-5ta cells revealed an increase in p53 Ser15 phosphorylation, an early event leading to cell senescence^25, 39^. Unexpectedly but interestingly, increased p53 Ser15 phosphorylation in senescent Cdk5rap2-depleted cells is not due to activation of the p53 kinases, ATM, Chk1, or Chk2^24^, but rather due to downregulation of WIP1 phosphatase expression. This concurs with previous reports that link cell senescence to WIP1-mediated p53 dephosphorylation at Ser15^28, 29^. While p53 phosphorylation at Ser15 was found to inhibit its interaction with the E3 ubiquitin ligase, Mdm2, preventing degradation and stabilizing p53^24^, we did not observe increased p53 level in Cdk5rap2-depleted cells, but only noted elevated p53 Ser15 phosphorylation. Since p53 Ser15 phosphorylation is required for p53 activation^25, 26^, WIP1-regulated p53 Ser15 phosphorylation and subsequent activation in Cdk5rap2-depleted cells likely account for senescence in these cells. Indeed, elevated level of p21^*CIP1*^, a major transcriptional target of p53^27^ that serves as a senescence biomarker^36^, was observed upon loss of Cdk5rap2. We note, however, that absence of γH2AX foci in Cdk5rap2-depleted cells indicates that p53-associated senescence that is linked to WIP1 downregulation in these cells is independent of DDR activation. This is consistent with the lack of DDR-associated activation of ATM, Chk1 or Chk2 in p53-associated cell senescence induced by loss of Cdk5rap2.

Our finding that knocking down WIP1 in Cdk5rap2-depleted cells causes increased p53 Ser15 phosphorylation and number of SA-β-gal positive cells, while ectopic expression of WIP1 in these cells reverses such phenotypes establishes a link between cellular senescence due to Cdk5rap2 loss and downregulation of WIP1. This is in accord with previous findings that WIP1^-/-^ mice exhibit aging phenotypes in their hematopoietic stem cells, including impaired repopulating activity^29^, and WIP1^-/-^ MEFs^29^ and mesenchymal stem cells^30^ exhibit premature senescence. In addition, reduced level of WIP1 in primary chondrocytes triggers the onset of senescence and loss of the chondrocyte phenotype, while overexpression of WIP1 retains their proliferative capacity and delays the onset of senescence^31^.

Our observation that Cdk5rap2 loss causes downregulation of WIP1 mRNA and protein levels together with the fact that the WIP1 promoter contains an NF-κB binding site through which NF-κB positively regulates WIP1 expression^32^, and β-catenin associates with NF-κB to induce the expression of NF-κB target genes^34^ led to our view that Cdk5rap2 loss reduces WIP1 level through downregulation of β-catenin. In addition, our notion that the level of β-catenin is regulated by GSK3β activity is based on the fact that GSK3β phosphorylates β-catenin, targeting β-catenin for ubiquitination-mediated degradation by the proteasome pathway^34^ and thus inhibiting translocation of β-catenin into the nucleus^44^. Our finding that Cdk5rap2 interacts with GSK3β and such interaction inhibits GSK3β activity explains how loss of Cdk5rap2 downregulates β-catenin level. The Harmonizome database^45^, and previous immunoprecipitation and mass spectrometry analyses^9^ support our finding that Cdk5rap2 binds to GSK3β. Additional evidence that reflects Cdk5rap2-GSK3β interaction is their colocalization in the centrosome, which duplicates in synchrony with the cell cycle. Indeed, although GSK3β, which promotes mitotic progression, is mainly distributed throughout the cytoplasm, a subpopulation of GSK3β localizes at the centrosome^46^. Recently, the phosphorylated form of GSK3β has been mapped to the centrosome^47^, where Cdk5rap2 exists^48, 49^ and regulates centriole engagement and cohesion during the cell cycle^43^. It remains to be determined whether disturbance of this Cdk5rap2 function contributes to senescence upon loss of Cdk5rap2. Nonetheless, since AKT (protein kinase B), which regulates centrosome composition and integrity during mitosis^50^, phosphorylates GSK3β at Ser9^51, 52^, it is possible that Cdk5rap2 binding to centrosome-recruited GSK3β facilitates the inhibitory phosphorylation of GSK3β Ser9 by AKT. Potentially, loss of Cdk5rap2 allows the release of unphosphorylated and active GSK3β from the centrosome, causing GSK3β phosphorylation of β-catenin, which is then ubiquitinated and degraded via the proteasome pathway. Reduced β-catenin level leads to reduced NF-κB-mediated WIP1 phosphatase expression, which results in sustained p53 Ser15 phosphorylation that promotes p53-mediated p21^*CIP1*^ expression, and ultimately induces cell senescence. Our finding that GSK3β inhibition restores β-catenin and WIP1 levels in cells lacking Cdk5rap2 establishes a link between increased GSK3β activity and decreased β-catenin and WIP1 levels in these cells. The fact that luciferase activity is reduced in WIP1 promoter-transfected cells depleted of Cdk5rap2 or β-catenin, and GSK3β inhibition restores luciferase activity in WIP1 promoter-transfected cells depleted of Cdk5rap2, further suggest that senescence caused by Cdk5rap2 loss is controlled by GSK3β through β-catenin regulation of the WIP1 promoter.

In summary, our studies reveal that loss of Cdk5rap2 triggers premature cell senescence via β-catenin-mediated downregulation of WIP1. Such senescence is associated with proliferation errors in both BJ-5ta cells and Cdk5rap2^*an/an*^ mice. As Cdk5rap2 loss-of-function has been linked to developmental defects, it is possible that premature cell senescence contributes to these defects. In this paper, we provide novel insight into the molecular mechanisms by which Cdk5rap2 loss-of-function causes premature cell senescence.

## Materials and Methods

### Materials

hTERT-immortalized BJ-5ta human foreskin fibroblast cells and adenovirus carrying H-Ras^*G12V*^ and GFP alone were obtained from Drs. Tara Beattie and Karl Riabowol, respectively, at the University of Calgary. HEK293 cells were from ATCC (Manassas, VA, USA). The Cdk5rap2 antibody (A300-554A) was from Bethyl Laboratories (Montgomery, TX, USA). p21^*CIP1*^ (sc-271610), p16^*INK4a*^ (sc-468), p53 (sc-393031), Ras (sc-35), histone H3 (sc-517576) and actin (sc-8432) antibodies, and protein A/G PLUS-agarose (sc-2003) were from Santa Cruz Biotech (Manassas, VA, USA). phosphoSer15-p53 (#9284), GSK3β (#12456), phosphoSer9-GSK3β (#5558), ATM (#92356), phosphoSer1981-ATM (#13050), Chk1 (#2360), phosphoSer345-Chk1 (#2348), Chk2 (#6334), phosphoThr68-Chk2 (#2197), β-catenin (#8480), GST (#2624) and Ki-67 (#9129) antibodies were from Cell Signaling (Danvers, MA, USA). WIP1 antibody (ab236515) was from Abcam (Cambridge, MA, USA). The HRP-conjugated anti-rabbit (#7074) and anti-mouse (#7076) secondary antibodies were from Cell Signaling (Danvers, MA, USA). Primers and siRNAs were synthesized at the University of Calgary Core DNA services or GenePharma, Shanghai, China. Goat anti-mouse or anti-rabbit secondary antibodies conjugated with Alexa-488 or −594 were from Invitrogen (Carlsbad, CA, USA). Myc-tagged Cdk5rap2 and Flag-tagged GSK3β expression vectors, and pReceiver-M02 carrying WIP1 were from GeneCopoeia (Rockville, USA). GSK3β protein (14-306) was from Upstate Biotech. (Lake Placid, NY, USA).

### Animals

The Cdk5rap2^*+/an*^ breeding pairs were purchased from JAX laboratory (Bar Harbor, ME, USA) and fed with commercial mice chow, provided water ad libitum and kept on a 12:12 hr light-dark cycle. Cdk5rap2^*+/+*^, Cdk5rap2^*+/an*^ and Cdk5rap2^*an/an*^ embryos were isolated from pregnant Cdk5rap2^*+/an*^ mice. All animal studies conformed to regulatory standards and were approved by the University of Calgary Health Sciences Animal Care Committee.

### Genotyping

Genomic DNAs were extracted from mice using the Extracta DNA prep kit (Quantabio, Edmonton, AB. Canada). Tissue samples were added to 50-100 µl of extraction buffer, incubated at 95°C for 30 min, and stabilized with an equal volume of stabilization buffer. DNA sample (1 µl) was added to 10 µl of Quantabio’s AccuStart II PCR SuperMix containing 100 nM of the appropriate DNA primers. The forward primers used were: GAAACCAGGGTGACA GGTACA (wt Cdk5rap2) and AGATGTCATGTCTAAAGCAATCACT (an Cdk5rap2). The reverse primer used was the same for the wt and an reactions: CCTTTGTCTTTCTGCCCTGA. Based on the expected size of each fragment, the wt *Cdk5rap2* allele corresponds to a 581 bp band whereas the mutant an *Cdk5rap2* allele corresponds to a 500 bp band. The reaction mixtures were input into a Bio-Rad thermocycler using PCR settings recommended by the Jackson Laboratory.

### Cell culture

BJ-5ta human diploid foreskin fibroblast cells were cultured in Eagle’s minimal essential medium (EMEM, Lonza) containing 10% fetal bovine serum (FBS, GIBCO), and 50 U/ml penicillin and 50 mg/ml streptomycin (Invitrogen, Carlsbad). HEK293 human embryonic kidney cells were cultured in Dulbecco’s modified Eagle medium (DMEM, Invitrogen), containing 10% FBS, 50 U/ml penicillin and 50 mg/ml streptomycin. Cells were maintained at 37°C in a 5% CO2 humidified incubator. After recovering from cryopreservation, BJ-5ta cells were used for up to 10 additional population doublings to maintains many characteristics of normal primary cells.

### Isolation of primary MEFs

Primary MEFs were isolated from E12.5 Cdk5rap2^*+/+*^, Cdk5rap2^*+/an*^ and Cdk5rap2^*an/an*^ embryos as described previously^53^. Briefly, embryos were washed with 1x PBS, decapitated and eviscerated then washed again with PBS. Embryos were minced using sterile forceps and placed in 3–5 ml of 0.05% trypsin-EDTA, pipetted up and down to get cells into suspension and incubated at 37°C for 5 min. Cell suspensions were transferred to tubes containing MEF medium (DMEM-high glucose supplemented with 10% FBS, 50 U/ml penicillin and 50 mg/ml streptomycin (Invitrogen, Carlsbad), and 2 mM GlutaMAX) then centrifuged at 1,000 rpm for 5 min. Cell pellets were resuspended in fresh media and plated in 10 cm cell culture dishes. Primary MEFs were cultured in DMEM supplemented with 10% FBS and 50 U/ml penicillin and 50 mg/ml streptomycin (Invitrogen, Carlsbad) under hypoxic condition (5% O2 and 5% CO2 incubator). All experiments were performed in passage P2-P7 MEFs.

### Plasmid/siRNA transfection and adenovirus infection

Cells cultured ∼18 hours and at about 60% confluency were transfected using Lipofectamine 2000 (Invitrogen) in serum-free medium, which was replaced with complete medium 5 hours post-transfection. Cells were harvested at different time points as indicated. siRNA target sequences are: control, CGUACGCGGAAUACUUCGAUU; Cdk5rap2 #1, GGACGUGUUGCUUCAGAAAUU; Cdk5rap2 #2, GAGUCAGCCUUCUGCUAAAUU; WIP1, CCAAUGAAGAUGAGUUAUAUU; GSK3β, AGGAGACCACGACCUGUUAAUU; β-catenin, CTCGGGATGTTCACAACCGAA. Adenovirus infection was carried out at a MOI of 50-100. Media was then replaced with EMEM containing 10% FBS 24 hours post infection and cells were incubated until the indicated time.

### RNA extraction and real-time qRT-PCR

Total RNA was extracted using TRIzol reagent (Invitrogen, Carlsbad, CA, USA) according to the manufacturer’s protocol, and transcribed into cDNA using high-capacity cDNA reverse transcription kit (Thermo Fisher, Waltham, MA). Real-time qRT-PCR was performed using power SYBR^®^ green PCR master mix (Thermo Fisher, Waltham, MA), and an Applied Biosystems 7500 real-time PCR machine using a standard protocol. The PCR conditions were 35 cycles at 94°C for 20 seconds, 60°C for 20 seconds and 72°C for 35 seconds. The primer sets used were: WIP1-F, GGGAGTGATGGACTTTGGAA; WIP1-R, CAAGATTGTCCATGCTCACC; GAPDH-F, GGAGCGAGATCCCTCCAAAAT; GAPDH-R, GGCTGTTGTCATACTTCTCATGG. GAPDH was used for normalization.

### Isolation of nuclear fraction

Lysates of BJ-5ta cells transfected with Cdk5rap2 siRNA or control siRNA were centrifugated at 800 rpm for 10 min. The resulting pellets were resuspended in cell lysis buffer containing 10 mM HEPES; pH 7.5, 10 mM KCl, 0.1 mM EDTA, 1 mM dithiothreitol (DTT), 0.5% Nonidet-40 (NP-40), 0.5 mM PMSF, and the protease inhibitor cocktail, then incubated on ice for 30 min with intermittent mixing, and centrifuged at 12,000 xg at 4°C for 15 min. The pellets were washed three times with cell lysis buffer, and resuspended and incubated in nuclear extraction buffer containing 20 mM HEPES (pH 7.5), 400 mM NaCl, 1 mM EDTA, 1 mM DTT, 1 mM PMSF and protease inhibitor cocktail for 30 min. Nuclear fractions were harvested by centrifugation at 12,000 xg for 15 min at 4°C.

### Western blot analysis

Cell lysates (50 μg) were resolved by SDS-PAGE, transferred to nitrocellulose membrane, and immunoblotted for Cdk5rap2, p21^CIP1^, p16^INK4a^, p53, phospho-p53, Ras, histone, GSK3β, phosphoSer9-GSK3β, ATM, phosphoSer1981-ATM, Chk1, phosphoSer345-Chk1, Chk2, phosphoThr68-Chk2, β-catenin, WIP1 and actin. Following incubation with HRP-conjugated anti-rabbit or anti-mouse secondary antibody, immunoreactive bands were detected using the ECL reagent (GE Healthcare, Little Chalfont, Buckinghamshire, UK). Western blot images were obtained using the ChemiDoc™ Imager (Bio-Rad) set at optimal exposure. No enhancements were performed.

### Immunofluorescence microscopy

Cells transfected with Cdk5rap2 or control siRNA on coverslips were fixed with 4% paraformaldehyde/PBS for 10 min, permeabilized using 0.1% Triton X-100/PBS for 10 min, then blocked in 2% BSA/PBS for 1 hour at room temperature. Cover slips incubated with the indicated primary antibody for 1 hour followed by 20 min incubation with secondary antibodies were washed with 1x PBS, counterstained with DAPI and mounted on glass slides using ProLong™ Diamond Antifade Mountant (P36961, Invitrogen, Carlsbad, CA, USA). Images were captured using a Zeiss Axiovert 200 microscope. BJ-5ta cells transfected with Cdk5rap2 siRNA #2 were analyzed for Ki-67 positive cells 3 days post-transfection.

### Senescence-associated β-galactosidase staining

Cells were fixed using 3% paraformaldehyde in PBS (pH 6.0) and stained with 1 mg/ml 5-bromo-4-chloro-indolyl-β-D-galactopyranoside (X-gal) solution containing 5 mM potassium ferrocyanide, 5 mM potassium ferricyanide, 150 mM NaCl, and 2 mM MgCl2 in PBS (pH 6.0) for 16-20 hrs at 37°C.

### Cell viability and cell cycle analyses

BJ-5ta cells transfected with Cdk5rap2 siRNA #2 or control siRNA were seeded into 96-well plates at 5,000 cells per well 24 hours post transfection, then analyzed for cell viability at days 0, 2 and 4 using Cell Counting Kit-8 (CCK-8, Dojindo). For analysis of cell cycle distribution, cells transfected with Cdk5rap2 siRNA #2 or control siRNA were fixed using cold 2% formaldehyde and stained with 7-AAD (5 μg/μl) then subjected to flow cytometry.

### Immunoprecipitation

HEK293 cells tranfected with Myc-tagged Cdk5rap2 or Flag-tagged GSK3β or both for 48 hrs were lysed in ice-cold lysis buffer containing 50 mM Tris/pH 8.0, 150 mM NaCl, 1% NP-40, 10 mM EDTA, 5% glycerol, 1 mM phenylmethylsulfonylflouride (PMSF), 10 µg/ml aprotinin, and 10 µg/ml leupeptin. Lysates were then clarified by centrifuged at 13,000 rpm for 25 min at 4°C. For immunoprecipitation under denaturing condition, 1% SDS was added in the lysis buffer. Lysates were pre-cleared by adding normal mouse or rabbit IgG + sepharose A/G beads and incubating at 4°C for 2 hrs followed by centrifugation at 14,000 rpm for 10 min. One-tenth of the lysates from each sample was retained and used to assess input or total cell lysate in western blots. Pre-cleared cell lysates were subjected to immunoprecipitation using Myc or Flag antibody conjugated to protein A/G agarose beads. After incubating the mixture overnight at 4°C, immunoprecipitates were washed with lysis buffer three times at 4°C.

### Measurement of GSK3β activity

GSK3β activity was measured using the GSK-3β activity assay kit (Sigma, ON, Canada) following the manufacturer’s protocol. Briefly, cells transfected with Cdk5rap2 or control siRNA were lysed in ice-cold lysis buffer containing 50 mM Tris/pH 8.0, 150 mM NaCl, 1 % NP-40, 10 mM EDTA, 5% glycerol, 1 mM phenylmethylsulfonylflouride (PMSF), 10 µg/ml aprotinin and 10 µg/ml leupeptin. Cell lysates (300 µg) were incubated with 2 µl of anti-GSK-3β and 30 µl EZview Red protein-G affinity gel beads at 4°C for 3 hrs. The beads were recovered by centrifugation at 8,000 xg for 30 sec and washing with 500 µl of ice-cold lysis buffer. The immunoprecipitates were mixed with 20 µl of reaction mixture containing 25 μCi γ^32^P-ATP and 5 µl of GSK-3β substrate solution, incubated at 37°C for 30 minutes, then spotted onto P81 phosphate cellulose membranes. Membranes were washed 4 times with 0.5% phosphoric acid and once with acetone. Counting of incorporated radioactivity was performed using a Beckman scintillation counter.

### Expression and purification of GST-Cdk5rap2

GST-Cdk5rap2 was cloned into pFastBac vectors from which baculovirus was generated according to the Bac-to-Bac^®^ baculovirus protein expression system (Thermo Fisher, Waltham, MA). Sf9 insect cells were infected with P2 baculovirus carrying GST-Cdk5rap2 for 24 hrs. GST-Cdk5rap2 was purified by affinity chromatography using a glutathione (GSH)-conjugated agarose column (Sigma-Aldrich, St. Louis, MO). Sf9 cell lysates expressing GST-Cdk5rap2 were incubated with GSH-agarose for 1 hour at 4° C and washed with 1x PBS containing 1% Triton X-100. Bound proteins were eluted with 10 mM reduced GSH in 50 mM Tris elution buffer (pH 8).

### GST pull down assay

GST-Cdk5rap2 (1 μmole) bound to glutathione-agarose beads was mixed with GSK3β (2 μg) at 4°C for 2 hrs. The pulled-down complex was washed 4 times with ice-cold GST lysis buffer (20 mM Tris-HCl (pH 8.0), 200 mM NaCl, 1 mM EDTA (pH 8.0), 0.5% NP-40, 2 µg/µl aprotinin, 1 µg/µl leupeptin, 0.7 µg/ml pepstatin and 25 µg/ml PMSF) by centrifugation at 2,500 rpm for 10 min and analyzed by SDS-PAGE and immunoblotting for GST and GSK3β.

### Generation of promoter constructs of wt WIP1 (pGL3-WIP1) and WIP1 with NF-κB binding site deletion (pGL3-WIP1-ΔκB)

Using genomic DNAs isolated from HEK293 cells as a template, the WIP1 promoter region was amplified by PCR using the primer set: ACATTTTCTTGAGCTGATTTTGCTT (WIP1-F1) and TCGGAGAAGACGCTCACTCC (WIP1-R1). The promoter region for WIP1-ΔκB was generated by PCR using two different sets of primers: ACATTTTCTTGAGCTGATTTTGCTT (WIP1-F1) and GTTTAAAAAGCACtta accgtcagct (WIP1-R2), and ACCGAGACTGTGCagctgacggttaaGTGCTT (WIP1-F2) and TCGGAGAAGACGC TCACTCC (WIP1-R1), with overlapping fragments (lowercase letters) in WIP1-R2 and WIP1-F2. Two PCR products were annealed and used as templates for subsequent 8 fusion PCR cycles^54^. PCR products were then purified using the GeneJET PCR Purification Kit (Thermo Fisher, Waltham, MA) and used as templates for PCR amplification using the WIP1-F1 and WIP1-R2 primer set. Generated WIP1 and WIP1-ΔκB promoter PCR products were cloned into pGL3-basic luciferase reporter vector (Promega, Madison, WI, USA) using XhoI and BglII restriction enzyme sites and designated as pGL3-WIP1 and pGL3-WIP1-ΔκB, respectively. Successful cloning was confirmed by DNA sequencing.

### Luciferase assay

To measure luciferase activity, HEK293 cells were co-transfected with the indicated siRNA, pGL3-WIP1 or pGL3-WIP1-ΔκB and pTK-Renilla luciferase vector using lipofectamine 2000 (Invitrogen, Carlsbad, CA, USA). Cell extracts were prepared 48 hours after transfection and luciferase activity was measured using the Dual-Luciferase Reporter assay system (Promega, Madison, WI, USA). Renilla luciferase served as an internal control for normalization.

### Statistical analysis

Student’s t-test (unpaired, two-sided) or one or two-way analysis of variance (ANOVA) was used. Significance was set at p<0.05.

## Acknowledgements

We thank Drs. Tara Beattie and Karl Riabowol at the University of Calgary for providing us BJ-5ta cells, and adenovirus carrying Ras V12 (Adeno-Ras V12) and control virus carrying GFP alone, respectively. We also thank Dr. Angela Kaindl at the Charité-Universitätsmedizin Berlin for providing a rabbit antibody against mouse Cdk5rap2. This work was supported in part by a grant from the NSERC (RGPIN/06270-2019) to KYL.

## Authorship contributions

XW performed most of the experiments and drafted the manuscript. PS and ZS performed the experiments for data presented in Figures 4 and 11, respectively. XG, JLR and KYL contributed to the analysis and interpretation of data and/or provided constructive comments on experimental design and/or contributed to the preparation and writing of the manuscript. JLR and KYL critically revised the manuscript for important intellectual content and wrote the final version of the manuscript.

## Supplementary Figure Legend

**Supplementary Figure 1.**
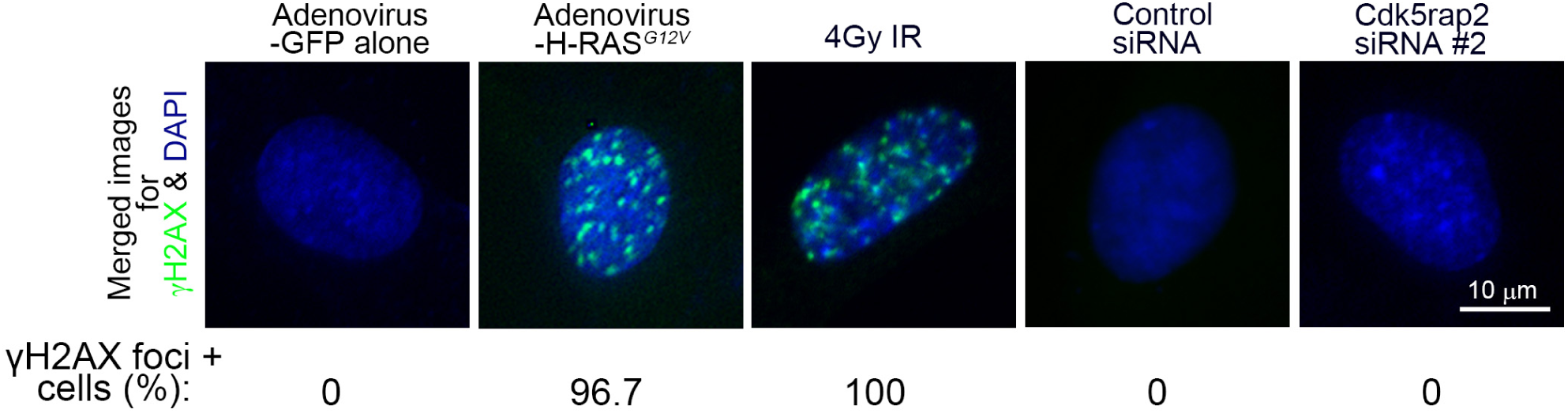
Cdk5rap2-depleted cells have no detectable γH2AX foci. BJ-5ta cells transfected with Cdk5rap2 siRNA #2 for 3 days were stained with DAPI and γH2AX antibody. Cells infected with adenovirus carrying H-RAS^*G12V*^ and cells treated with 4Gy IR were used as positive controls for γH2AX detection. Merged images of γH2AX and DAPI staining are shown. Representative images are from one of three independent experiments showing similar staining patterns. The indicated percentage (%) of γH2AX positive cells was assessed in ∼200 cells per treatment group in each of the 3 independent experiments (n=3).

**Supplementary Figure 2.**
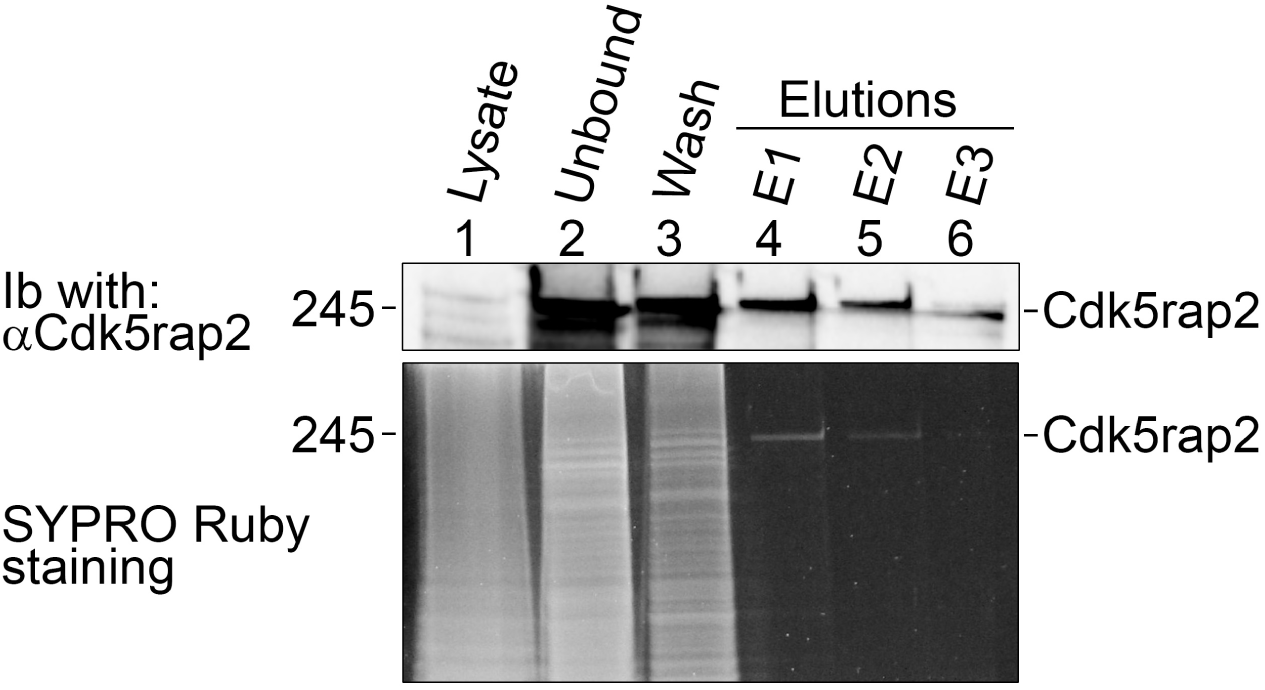
Purification of GST-Cdk5rap2. Lysate of Sf9 insect cells expressing GST-tagged Cdk5rap2 was subjected to affinity purification using a glutathione-agarose column. After collecting the flow-through, the column was washed until no more protein was detected in the wash. Bound proteins were eluted from the column with elution buffer containing 10 mM reduced glutathione. Three elution fractions (E1-E3) were collected and subjected to SDS-PAGE and immunoblotting for Cdk5rap2 (upper panel). SYPRO Ruby staining (lower panel) was performed to assess protein loading and GST-Cdk5rap2 purification.

**Supplementary Figure 3.**
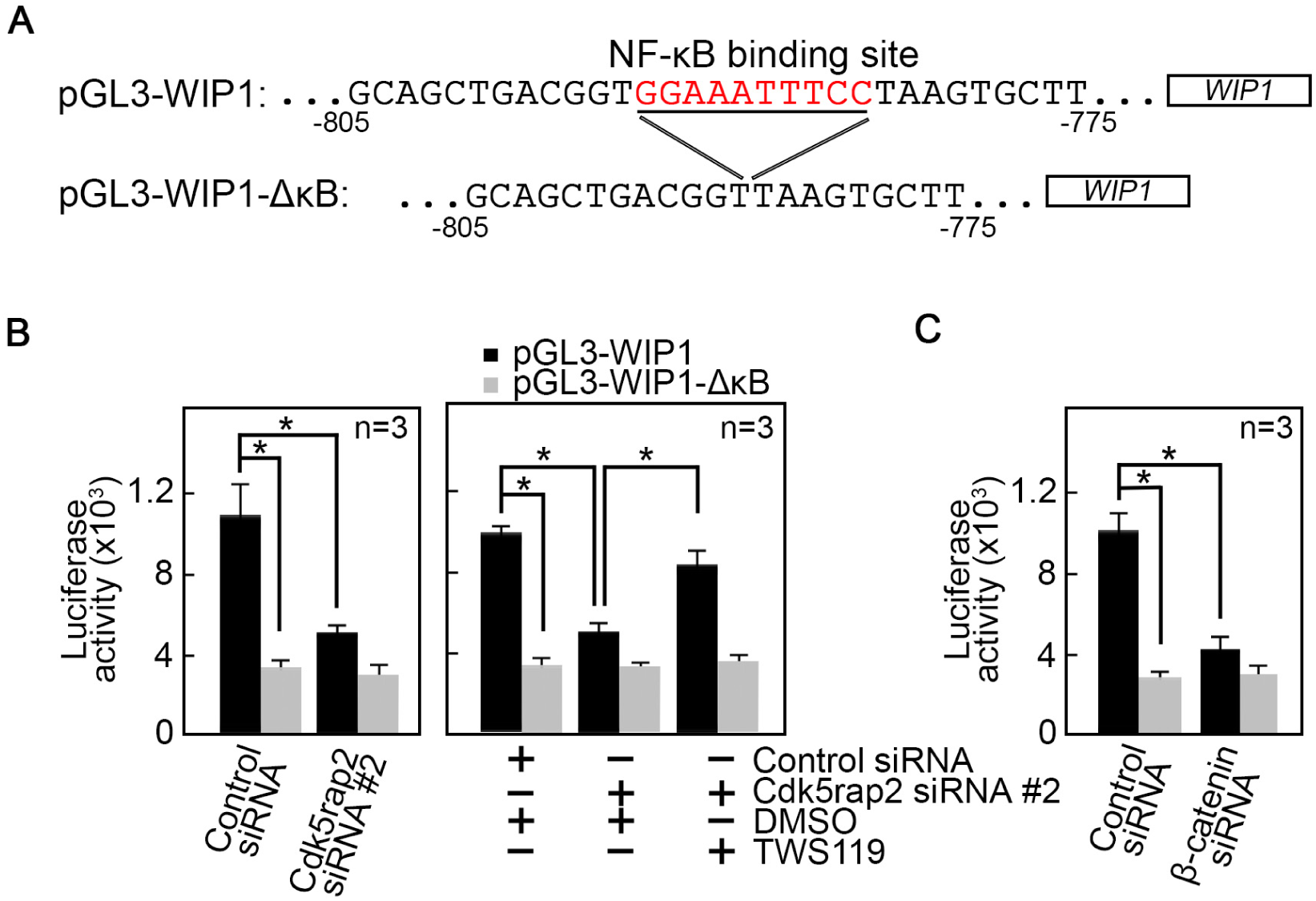
Cdk5rap2 regulates WIP1 promoter activity via β-catenin. A. Sequence of pGL3 luciferase reporter vector carrying WIP1 or WIP1-ΔκB promoter. B and C. Loss of Cdk5rap2 (B, left and right panels) or β-catenin (C) inhibits WIP1 promoter activity as measured by luciferase reporter activity but inhibition of GSK3β with TWS119 in Cdk5rap2-depleted cells restores WIP1 promoter activity (B, right panel). Data represent means ± SD from three (n=3) independent experiments. *p<0.05.

